# Disentangling genomic imprinting from indirect parental genetic effects on the human methylome

**DOI:** 10.64898/2026.11.18.767489

**Authors:** Fernanda Adame Galeano, Ilse Krätschmer, Anne Richmond, Caroline Hayward, Riccardo E. Marioni, Beatriz Vicoso, Matthew R. Robinson

## Abstract

Imprinted genes regulate development and metabolism and are implicated in both rare developmental syndromes and common diseases. Genomic imprinting is conventionally inferred from parent-of-origin effects (POEs), identified as differences between heterozygotes who inherited an allele from their mother and those who inherited it from their father. However, such differences can also be caused by indirect parental genetic effects, i.e. the fact that the genotype of a parent can be associated with behavioral or environmental changes that affect the phenotype of their offspring. No study of the human methylome has so far accounted for this confounding effect when inferring imprinted loci. Here, we jointly estimate direct, maternal, paternal and parent–of–origin effects at 850,055 methylation probes from whole blood in 2,438 parent– offspring trios from Generation Scotland. Of 5,435 probes with apparent POE signal, 4,974 (91.5%) fall below threshold once parental genotypes are included, suggesting that the vast majority are not attributable to imprinting alone. We treat the 281 remaining probes, 17 on the X chromosome and 42 not previously reported, as a conservative, high–confidence set, map them to 1,118 SNP loci linked to gene expression and protein abundance, and identify 27 probes where imprinting and indirect parental genetic effects both contribute. In a hold–out set of relatives, 250 of the 281 show familial resemblance consistent with a combination of imprinting and parental effects. Our results highlight the importance of accounting for indirect parental genetic effects when interpreting putative imprinting regions.

**Significance:** Some genes behave differently depending on which parent they were inherited from, an effect called genomic imprinting that shapes development, metabolism and disease risk. Imprinting is detected by comparing children who inherited the same DNA variant from their mother versus their father. But parents also influence their children through the environment they provide, and these two effects are statistically entangled: each can look exactly like the other unless the parents’ own DNA is controlled for. We separate them using more than 2,400 families with genetic and epigenetic data on both parents and the child. Nine in ten apparent imprinting signals do not survive conditioning on parental genotypes, highlighting the importance of accounting for indirect parental genetic effects.

---

Parent–of–origin effects (POEs) occur when the expression of an allele depends on the sex of the parent from whom it was inherited. Genomic imprinting is considered the primary mechanism underlying POEs and is characterized by differences in DNA methylation between maternally and paternally inherited sequences (imprinted differentially methylated regions, iDMRs) (1). iDMRs are either established in gametes and escape the epigenetic reprogramming that follows fertilization, or are established after fertilization and persist through adulthood (2). iDMRs regulate the expression of either maternal or paternal alleles at nearby genes or gene clusters, which can be observed as a difference between heterozygous individuals who inherited their alternative allele from their father and those who inherited it from their mother (reciprocal heterozygotes) (1). Disruption of normal imprinting patterns is associated with several developmental disorders, and growing evidence links imprinting to susceptibility to complex diseases (3–7). Identifying where POEs occur in the human genome and distinguishing them from effects that merely resemble them, such as indirect genetic effects described in detail below, is a key challenge in human genetics.

DNA methylation is dynamic across the life course and can be influenced by developmental processes, environmental exposures, and lifestyle factors (8). During gestation, the fetal epigenome undergoes remodeling, making it uniquely vulnerable to maternal and environmental inputs, which may influence a child’s lifelong health trajectory (9). Parents differ in the prenatal and postnatal environments they provide, creating differences in methylation among children. If these differences reflect heritable parental traits, then indirect parental genetic effects may occur, where differences among parents in their genotype influence the phenotype of children beyond the transmission of alleles from parents to children (10).

For example, indirect maternal genetic effects may occur when a mother’s genotype influences her metabolic or behavioral traits, which in turn modify the fetal environment, creating lasting epigenetic changes in the offspring. Despite their potential importance, indirect parental genetic effects on DNA methylation are only beginning to be systematically characterized (10). In particular, no study to date has separated the two potential paths through which parental DNA can influence offspring methylation: (i) directly, by influencing the establishment of sex–specific methylation marks during gametogenesis (sperm and oocyte development) that underlie iDMRs and (ii) indirectly, through parental genotypes influencing parental phenotypes and thus intrauterine and early–life environments. Thus, distinguishing between the two mechanisms is critical, as true imprinting regions may be masked by indirect parental genetic effects, while previously identified POE-associated loci may reflect indirect parental genetic effects rather than imprinting.

Existing methods to test whether the effects of specific alleles on traits of interest differ between reciprocal heterozygotes have a key limitation: there is a correlation between parental genotype and the probability of inheriting an allele. This is a consequence of homozygous parents being able to produce only one kind of reciprocal heterozygote. It makes differences among reciprocal heterozygotes impossible to separate from differences due to parental genotypes, without jointly modeling both POEs and indirect genetic effects (i.e. estimating POE contributions after conditioning on parental genotypes and vice versa) (11, 12). Without this distinction, genuine imprinting regions may be missed, and previously detected POE loci associated with common complex disease (13), gene expression (14) and plasma protein levels (13), as well as currently known iDMRs (15–17) may not be caused by imprinting, but rather reflect an association with parental physiology, behavior or the environment.

Here, we identify differentially methylated regions as those for which the methylation patterns in children differ between reciprocal heterozygotes for single nucleotide polymorphisms (SNPs) within the same DNA region (those in ‘cis’), while controlling for potential indirect parental genetic effects. We use data from the largest within–family methylation study conducted to date, Generation Scotland, where methylation levels are quantified for 18,843 participants. We discover 281 methylation probes across the whole genome that are differentially methylated between maternal and paternal chromosomes in whole blood, 89% of which replicate in a hold–out set of related individuals when tested for patterns of familial resemblance consistent with imprinting. The majority of our high–confidence POE–dependent probes overlap with loci previously implicated in imprinting, supporting the robustness of our approach. For other previously reported loci, a primary contribution of indirect parental genetic effects could not be excluded. We further identify 42 novel POE loci with associated genes extending to known imprinting pathways. The POE–associated SNPs overlap with existing functional genomics datasets: a substantial proportion coincide with loci associated with gene expression, protein abundance and phenotypes, highlighting the importance of accounting for indirect parental genetic effects when defining the landscape of human imprinting.

## Results

### Study overview

We distinguish indirect parental genetic effects and POEs using a custom form of multivariable linear regression, where the response variable is the children’s methylation level at a given DNA position (as measured through a probe, the term we use to refer to these methylation levels). The methylation probes are regressed on a design matrix where for a given SNP *j* = 1, …, *p* for *p* SNPs, we jointly estimate four regression coefficients ***β***_*j*_ = (*β*_*Cj*_, *β*_*Mj*_, *β*_*F j*_, *β*_*Ij*_)^*T*^ for: (i) the child’s genotype (direct effect, *β*_*C*_); (ii) the mother’s genotype (indirect maternal genetic effect, *β*_*M*_ ); (iii) the father’s genotype (indirect paternal effect, *β*_*F*_ ); and (iv) the parent– of–origin indicator (POE, *β*_*I*_ ), determined by phasing the genotypes (see Figure 1). We jointly analyze all SNPs on the chromosome on which the probe is located, to estimate the effects of each SNP conditionally on all other SNPs of the same chromosome. Previously, this model has been applied to show that genetic effects for height, body mass index and educational test scores contain a small but significantly non–zero contribution from both parental genetic effects and POEs (18), but it has not been applied to give a mechanistic understanding of imprinting.

**Fig 1.**
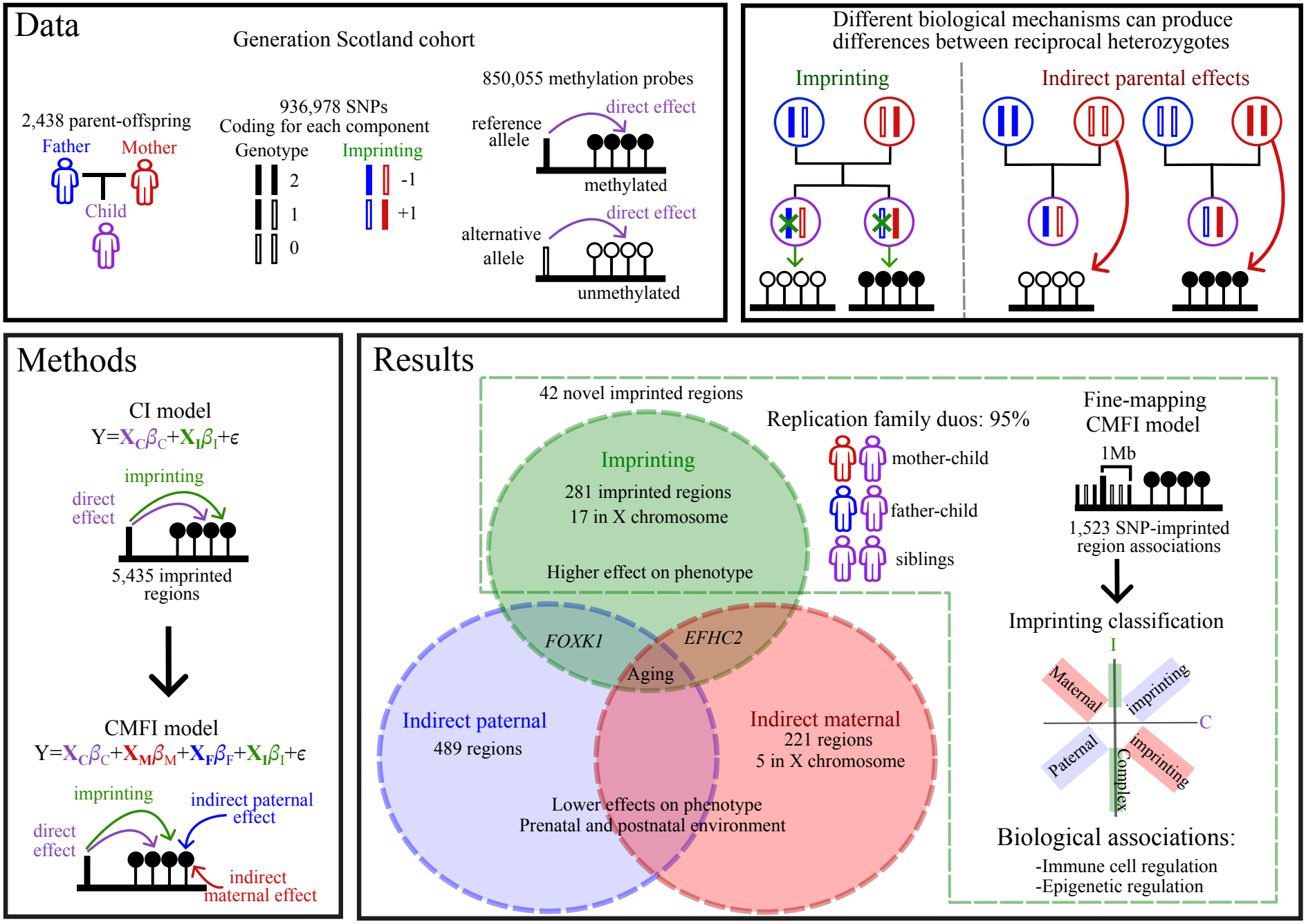
Study overview. The dataset used, examples of imprinting and indirect parental genetic effects for reciprocal heterozygotes, the two–phase framework used to identify POE–dependent probes and probes with significant contributions from indirect maternal and paternal genetic effects, and overview of the results, including replication, fine–mapping, classification of imprinting patterns and downstream biological associations for POE–dependent probes. See text for more details.

We apply this iterative Bayesian method to data from Generation Scotland (GS), which contains methylation values for 18,843 participants at 831,349 autosomal and 18,706 non–pseudoautosomal X chromosome loci within the genome using an Infinium MethylationEPIC BeadChip in whole blood. Fully phased imputed data of 11,358,693 autosomal and 16,104 X chromosome SNPs are also available, of which we use a subset of 931,607 common autosomal and 5,371 X chromosome SNPs for the identification of POE–dependent probes. We conduct extensive simulation studies to *(i)* validate the application of our statistical approach to these data; *(ii)* to ensure that POEs can be reliably distinguished from indirect parental genetic effects; and *(iii)* to demonstrate that indirect parental genetic effects can be misattributed to POEs (or *vice versa*, POEs being misattributed to indirect effects). The simulation studies are presented in Supplementary Figures S1–S2 and discussed in the Supplementary Text.

### Identification of genomic regions with differential methylation between maternal and paternal chromosomes

Using a subset of phased genotype data with strict quality controls (931,607 common autosomal SNPs in 2,438 child–mother–father trios, and 5,371 X chromosome SNPs in the 1,182 of those trios with a daughter; see Materials and Methods), we discover 281 methylation probes across the whole genome that show POE–dependent methylation, 17 of which are located on the X chromosome (Figure 2c).

**Fig 2.**
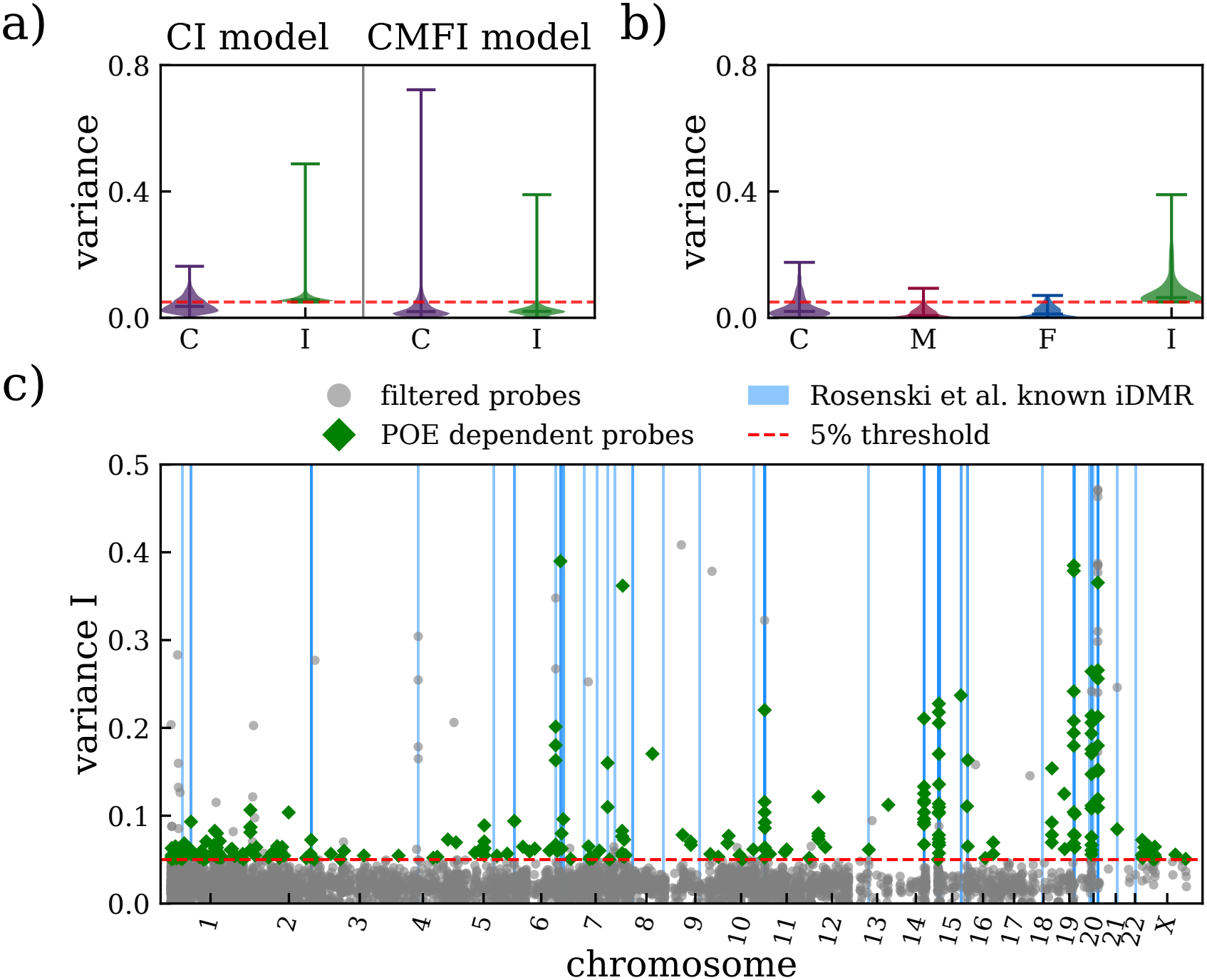
Estimated variances for POE–dependent probes. (a) Distribution of variances due to direct genetic (C) and imprinting effects/POEs (I), estimated using both the *CI* model and the *CMFI* model (see Materials and Methods), for 5,435 probes initially identified as having significant variance attributed to POEs (*V*_*I*_ *>* 5%) using the *CI* model. The variance attributed to POEs is substantially reduced after accounting for indirect parental genetic effects (*CMFI* model), highlighting their importance for the accurate identification of true imprinting signals. The red line indicates the 5% variance threshold. (b) Distribution of variances due to direct genetic (C), indirect maternal (M), indirect paternal (F) and imprinting effects (I) estimated using the *CMFI* model, for the 281 identified POE–dependent probes. (c) Variance due to POEs as a function of genomic location for probes initially identified under the *CI* model, which are filtered after running the *CMFI* model (grey circles), and for the final set of POE–dependent probes identified (green diamonds). Vertical blue lines indicate the positions of known imprinted differentially methylated regions (iDMRs) from Rosenski *et al*. (15).

We first scan all 850,055 probes using a model that includes only direct genetic effects and POEs (*CI* model) and find 5,435 methylation sites that show significant evidence for imprinting effects, where we recover *>* 5% variance attributable to the *I* component, *V*_*I*_ . We then re–run the analysis for these probes adjusting for parental genotypes using a model that includes genotype effects of the child, the parents and POEs (*CMFI* model) and find that the estimated variance due to POEs decreases sub-stantially for most probes, with 4,974 (91.5%) falling below the 5% threshold (Figure 2a). After additionally filtering for uncertainty in the estimated variance components, we obtain 281 probes where the variance attributable to imprinting remains at *>* 5% after conditioning on parental genotypes. We conservatively infer these probes to be POE–dependent. Imprinting is not the only source of genetic influence at these loci: direct effects (*C* component, *V*_*C*_) explain more than 5% of the variance for 61 probes, while indirect paternal (*F* component, *V*_*F*_ ) and indirect maternal (*M* component, *V*_*M*_ ) genetic effects contribute significantly to the variance of 19 and 11 probes (including 3 with all significant effects), respectively, indicating that even at methylation sites where POEs play a major role, other sources of genetic variation frequently combine to meaningfully influence the phenotype. Figure 2b shows the distributions of the estimated variance due to the different effects for the 281 identified POE–dependent probes. A list of the POE–dependent probes and their estimated effects is provided in Supplementary Dataset S1.

### Fine–mapping SNPs within imprinted regions

For the identified set of POE–dependent probes, we identify associated SNPs with a posterior inclusion probability (PIP *≥*0.95) and an estimated effect 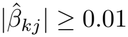, with *k* = 1, … *K* for *K* components. To fine–map the potential causal variants underlying these signals, we define genomic regions of 1 Mb on each side of the associated SNP. Each associated SNP is treated independently; thus, for each identified POE–dependent probe, all imputed SNPs within a 2 Mb window of the originally identified SNP are extracted, excluding the *r*^2^ filter, and reanalyzed using the *CMFI* model. We define credible sets of SNPs, as SNPs with pairwise *r*^2^ *≥*0.5 and retain those with a PIP *≥*0.95.

Because each originally associated SNP is fine–mapped independently, the same fine–mapped variant can be recovered from more than one regression model using slightly different fine–mapping windows. This yields a total of 2,393 fine–mapped SNP–probe associations, of which 1,523 associations are unique between 1,118 SNPs and 119 POE–dependent probes. This represents 42% of the POE signals having a fine-mapped variant association that passes the PIP *≥*0.95 threshold (Figure 3 and Supplementary Dataset S2). In many cases, multiple SNPs are associated with the same probe, suggesting polygenic contributions (Figure 3c). We also observe multiple instances of pleiotropy, as many SNPs are associated with more than one POE–dependent probe (Figure 3d).

**Fig 3.**
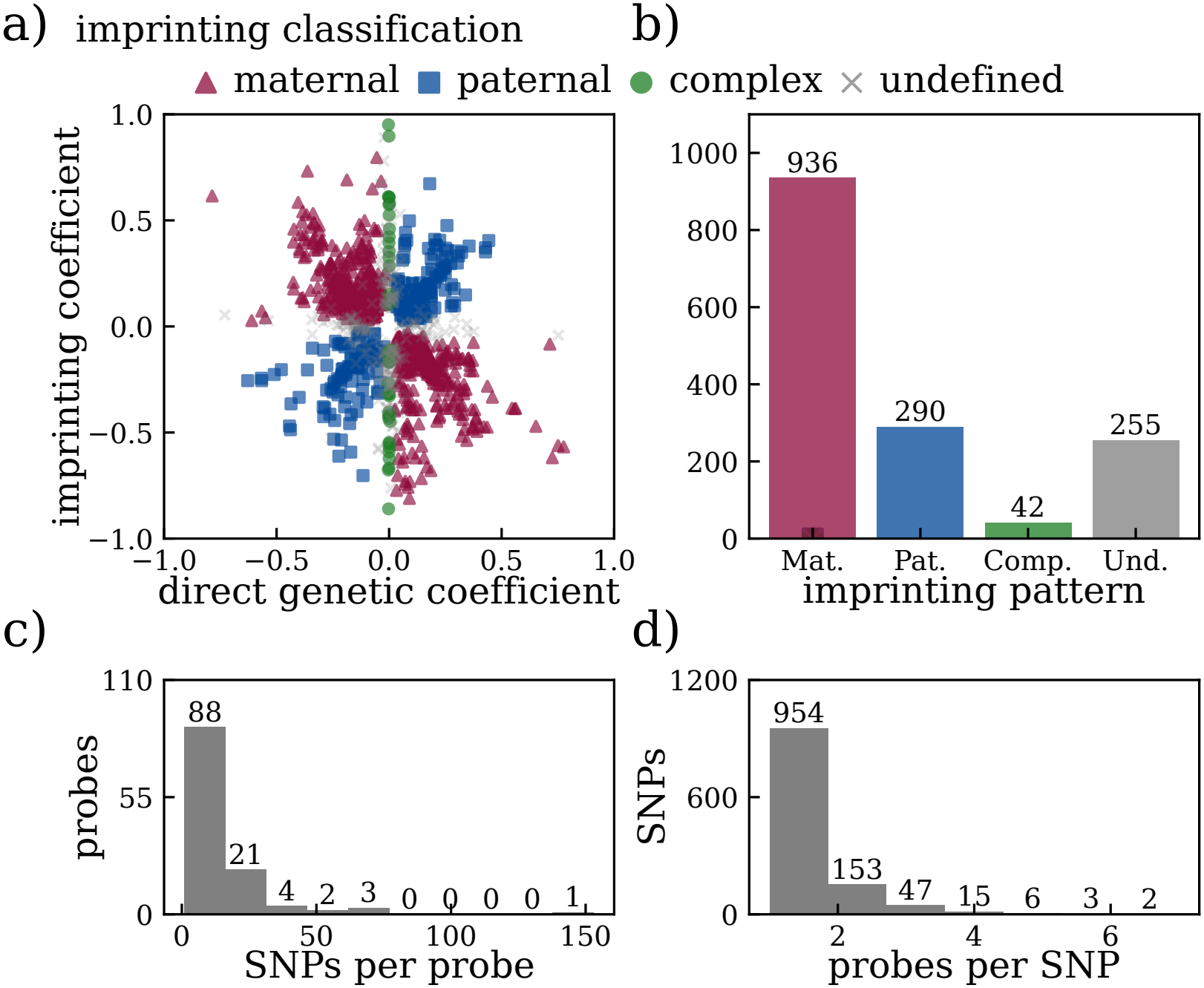
Characterization of fine–mapped SNPs associated with POE–dependent probes. (a) Estimated direct genetic and imprinting effect for all fine–mapped SNPs, used to classify imprinting patterns. (b) Number of SNP-probe associations allocated to each imprinting pattern. (c) Histogram of the number of fine–mapped SNPs identified per probe, consistent with polygenic contributions to methylation variance. (d) Histogram of the number of POE–dependent probes identified for each fine–mapped SNP, indicative of pleiotropy.

### Classification of imprinting patterns

Using the SNP–probe associations obtained with fine–mapping, we can infer paternal and maternal imprinting based on the signs of the SNP regression coefficients *β*_*C*_ and *β*_*I*_ (Figure 3a,b).

The signs of *β*_*C*_ and *β*_*I*_ together identify the parental copy through which the variant acts. When *β*_*C*_ and *β*_*I*_ share a sign (*β*_*C*_*β*_*I*_ *>* 0), the effect of the alternative allele on methylation appears only when that allele is maternally inherited, implying that the paternally inherited copy is silenced (paternal imprinting). Opposite signs (*β*_*C*_*β*_*I*_ *<* 0) imply the reverse: maternal imprinting, with expression from the paternal allele. This classification assumes that the unresponsive copy is the transcriptionally silenced one and, thus, that higher methylation results in lower expression. SNPs where *β*_*C*_ is indistinguishable from zero are classified as complex and represent patterns influenced by dominance effects such as bipolar, over–, and under–dominance. In these patterns, the two homozygotes do not differ in their average methylation levels, while methylation differences remain between the reciprocal heterozygotes (19). SNPs where we are unable to resolve the observed pattern are classified as undefined. For more details, see Materials and Methods.

The top panels of Figure 4 show examples of simulated methylation patterns, assuming paternal imprinting (maternal allele expression) (left) and indirect maternal genetic effects without imprinting (right). The methylation patterns of all children for the two different scenarios look similar. However, we can differentiate between imprinting and indirect effects when looking at the methylation of reciprocal heterozygotes from heterozygous parents (top right of each example) and at differences between siblings (bottom right of each example). When indirect effects exist, we expect similar methylation values for reciprocal heterozygotes of the parent causing the effect, and no differences among any type of sibling pair, since siblings share both parents and within-pair differences cancel indirect parental effects.

**Fig 4.**
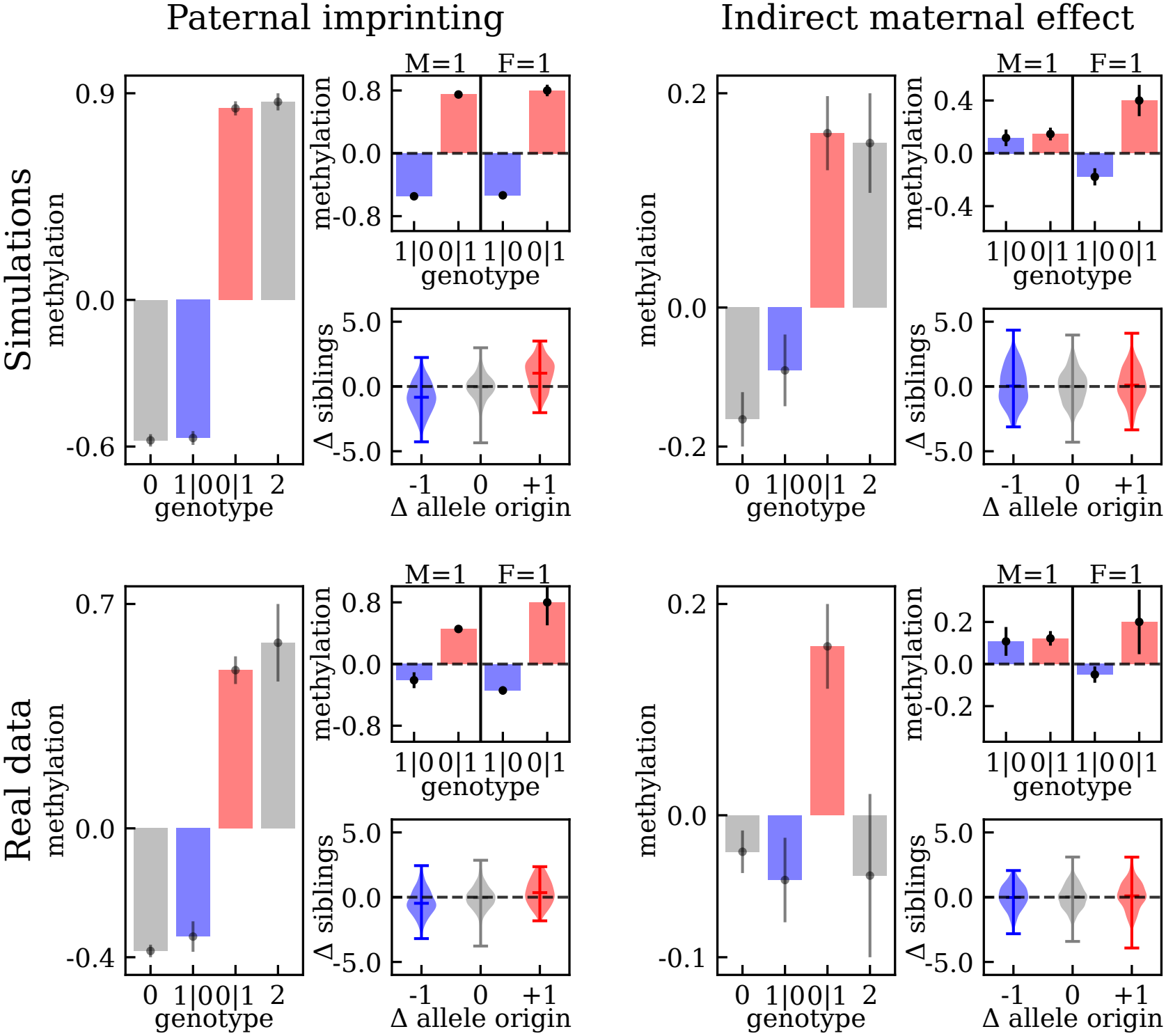
Examples of paternal imprinting and indirect maternal genetic effects. Methylation patterns reflecting paternal imprinting (left) and indirect maternal genetic effects (right), shown for simulated data (top) and for real data (bottom): SNP rs72698721 acting on probe cg04291079, associated with *MEG3* (bottom left), and probe cg07531550, which presents indirect maternal genetic effects (bottom right). For each example, three plots are shown: (i) average methylation of all children adjusted for the selected SNP and stratified by genotype, where differences between reciprocal heterozygotes (shown in blue and red) are typically interpreted as evidence of imprinting; (ii) average methylation values of reciprocal heterozygote children from heterozygous parents adjusted for the maternal *M* or paternal *F* indirect component of the SNP, where similar measurements for both reciprocal heterozygotes from either heterozygous mothers or fathers point towards indirect parental genetic effects; and (iii) violin plots of differences in methylation of sibling pairs adjusted for the imprinting *I* component, stratified according to parental origin of the alternative allele, where a slope between different sibling pairs indicates an imprinting effect. The error bars in (i) and (ii) represent the standard deviations.

The bottom panels of Figure 4 show real data examples for one associated SNP of probes cg04291079 (left) and cg07531550 (right). Probe cg04291079 at *MEG3* has a clear paternal imprinting pattern across 22 associated SNPs. In contrast, the variance attributed to POEs estimated for probe cg07531550 at *FRG1* under the *CI* model is 9.22% (95% posterior credible interval 9.06–9.39%), which reduces to 2.26% (95% posterior credible interval 2.12–2.40%) under the *CMFI* model. At the same time, the variance due to indirect maternal genetic effects rises to 9.29% (95% posterior credible interval 9.02– 9.55%), identifying this as an indirect maternal genetic effect rather than true imprinting (Figure 4).

The imprinting patterns that we infer show strong concordance with those previously reported in the literature. We identify several SNPs with maternal imprinting patterns that are linked to well known maternally imprinted genes, such as *DLGAP2* (60 SNPs), *BLCAP* (53 SNPs), and *DIRAS3* (33 SNPs). Likewise, we find many SNPs with paternal imprinting patterns that are associated with well known paternally imprinted genes, such as *MEG3* (22 SNPs, representing 84.6% of all defined imprinting patterns associated with this gene), *PWRN3* (9 SNPs) and *NUDT12* (5 SNPs). However, some loci require additional interpretation. For example, SNP rs7481173, associated with probe cg02425416, is linked to *INS–IGF2* and *IGF2*, which are typically described as paternally expressed. In our study, however, all SNPs with a defined imprinting pattern fine–mapped from the originally identified rs7481173 acting on cg02425416 are classified as complex. Upon closer inspection, a clear bipolar imprinting pattern can be observed, as shown in Supplementary Figure S3. Another example is *RB1*, which is classified as maternally imprinted for 25 associated SNPs, despite being known to be preferentially expressed from the maternal allele. This is consistent with its regulatory mechanism, where the unmethylated paternal allele acts as a promoter of an alternative transcript that biases expression towards the maternal allele (20). Furthermore, *MEG8* is classified as maternally imprinted across 17 SNPs, consistent with the tissue–specific switch to maternal imprinting in whole blood reported by Zhou *et al*. (21) (Supplementary Figure S4).

### 42 POE–dependent probes not previously reported

We compare our results to previous family based studies: (i) parental allele specific methylation and known imprinted differentially methylated regions from Rosenski *et al*. (15); (ii) known imprinted regions and POE–dependent probes reported by Cuéllar-Partida *et al*. (17); and (iii) POE– influenced probes identified in a previous analysis in the Generation Scotland cohort by Zeng *et al*. (16).

We find 42 POE–dependent probes that have not been reported previously; we consider this number a conservative estimate of novelty. We also replicate 81% of the known imprinted regions reviewed by Rosenski *et al*. using a 2 Mb location window (Figure 2c and 5a).

Because Zeng *et al*. used the same cohort, we separately compare our results against only the two studies that do not use Generation Scotland data (Rosenski *et al*. and Cuéllar-Partida *et al*.), to assess novelty independent of cohort overlap (Figure 5). This comparison identifies 205 overlapping probes, 76 probes not found in either study (not identified in non–GS studies), and 19 probes reported in these studies that we did not identify as POE–dependent. The 76 probes novel relative to non–GS studies exhibit significantly higher variances attributable to indirect parental genetic effects compared to over-lapping probes (maternal: *P* = 1.75 *×* 10^*−*21^, paternal:*P* = 3.14 *×*10^*−*17^) and probes not identified in our study (maternal: *P* = 1.33 *×*10^*−*18^, paternal: *P* = 5.33*×* 10^*−*19^). In contrast, their variances due to imprinting effects are lower (*P* = 5.64*×* 10^*−*13^ and *P* = 2.98*×* 10^*−*6^). These findings suggest that our approach detects imprinting signals even when these are smaller and in the presence of indirect parental genetic contributions.

**Fig 5.**
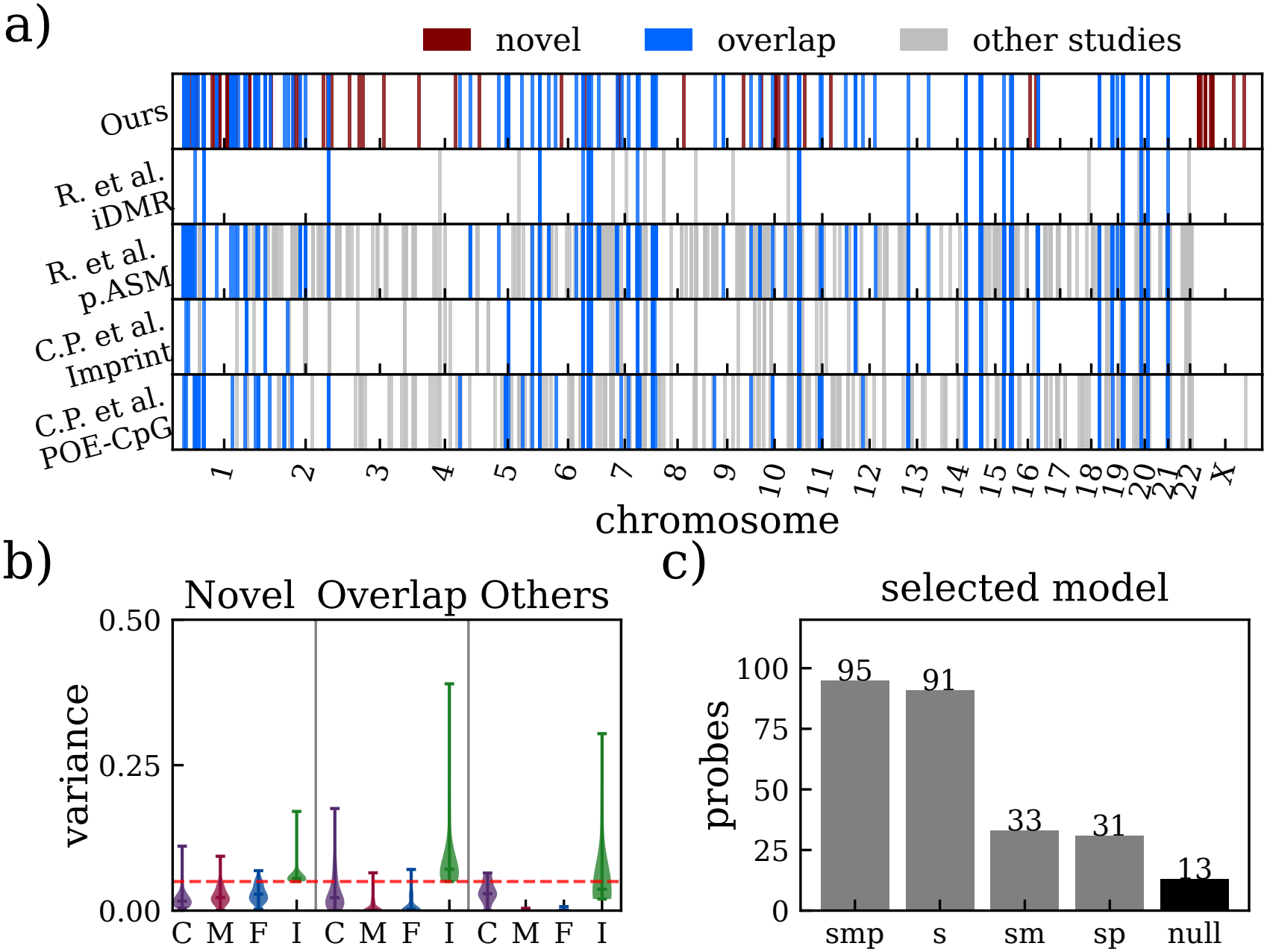
Comparison with previous family–based studies not using GS data and replication with an independent set. (a) Genomic locations of POE–dependent probes identified in this study (Ours), known imprinted differentially methylated regions from Rosenski *et al*. (15) (R. et al. iDMR), parental allele–specific methylation regions from Rosenski *et al*. (15) (R. et al. p.ASM), imprinted regions from Cuellar–Partida *et al*. (17) (C.P. et al. Imprint), and POE–CpG regions from Cuellar–Partida *et al*. (17) (C.P. et al. POE–CpG). Overlapping probes (blue) are found in our study and in at least one external dataset. Regions present only in external datasets (gray) are classified as “other studies”. (b) Violin plots for the distributions of variances due to direct (C), indirect maternal (M), indirect paternal (F) and imprinting effects (I) estimated under the *CMFI* model for 76 novel POE–dependent probes relative to non–GS studies identified in our study, 205 overlapping probes, and 19 probes only found in previous studies, whose estimates did not pass the uncertainty threshold or whose variance attributed to POEs did not pass the 5% threshold, indicated by a red dashed line. (c) Replication of POE–dependent probes identified in our study using a pedigree–based variance component framework adapted from Zeng *et al*. (16): models with different relatedness matrices representing different family resemblance structures are tested against the null model not including relatedness matrices. The number of probes is shown for each selected model, where smp represent the sibling–mother–father model; s the sibling model; sm the sibling and mother model; sp the sibling and father model; and null the null model of no increase in familial sharing of methylation values.

### Replication shows familial resemblance consistent with imprinting

Using a hold–out set of the Generation Scotland data containing 3,854 parent–offspring duos and 4,856 sibling pairs from 13,564 individuals who are not part of any of our parent–offspring trios, we replicate our findings by extending the approach described in Zeng *et al*. (see Materials and Methods). We replicate 250 out of 264 (94.7%) autosomal POE–dependent probes that we tested, with only one autosomal probe not converging. POE– associated loci that were not recovered in this additional dataset did not differ significantly in estimated variance components or genomic location compared to those that were, suggesting that these variables do not explain the replication rate. This approach is based on the assumption that a model with additional familial relationships is expected to perform better than a null model, with replication testing restricted to autosomal probes. The majority of the replicated probes are identified both in the model including maternal, paternal, and sibling relationships and in the model including only sibling relationships. It should also be noted that in this model, an improvement in model fit to the data may not be specific to imprinting, as it can also arise from other sources of family resemblance, such as indirect parental genetic effects, shared household or prenatal environment, or non–additive genetic variance. Thus, our findings are consistent with the idea that the imprinted regions we identified are influenced by multiple sources of genetic and familial effects, including indirect parental genetic effects (Figure 5c).

### Associated genes and potential functional consequences of POE–dependent methylation

We find 176 unique gene identifiers linked to POE–dependent probes and their fine–mapped SNPs (see Materials and Methods, Supplementary Dataset S3). Of these, 133 are detected neither in the POE studies mentioned above nor in the parent–of–origin eQTLs from Guan *et al*. (14), greatly expanding the known set of genes potentially affected by POE.

We search for known or predicted interactions among our set of POE-associated genes using STRING DB (22). The resulting network includes a functional cluster containing many known imprinted genes, such as *PEG3, KCNQ1, GNAS* and *IGF2*. Importantly, 39 of our novel POE-associated genes are also connected to this network (Figure 6). These include *HDAC1, RING1, BCOR, FOXK1,FABP7* and *EFHC2*, which are involved in development and/or epigenetic regulation, and are promising candidates for future investigation. A functional enrichment analysis highlights metabolic disease, birth weight, and body weight (False discovery rate, FDR = 1.8 *×*10^*−*4^, 1.4 *×*10^*−*4^, and 6 *×*10^*−*4^ respectively), consistent with the established roles of imprinted genes (Supplementary Figure S6).

**Fig 6.**
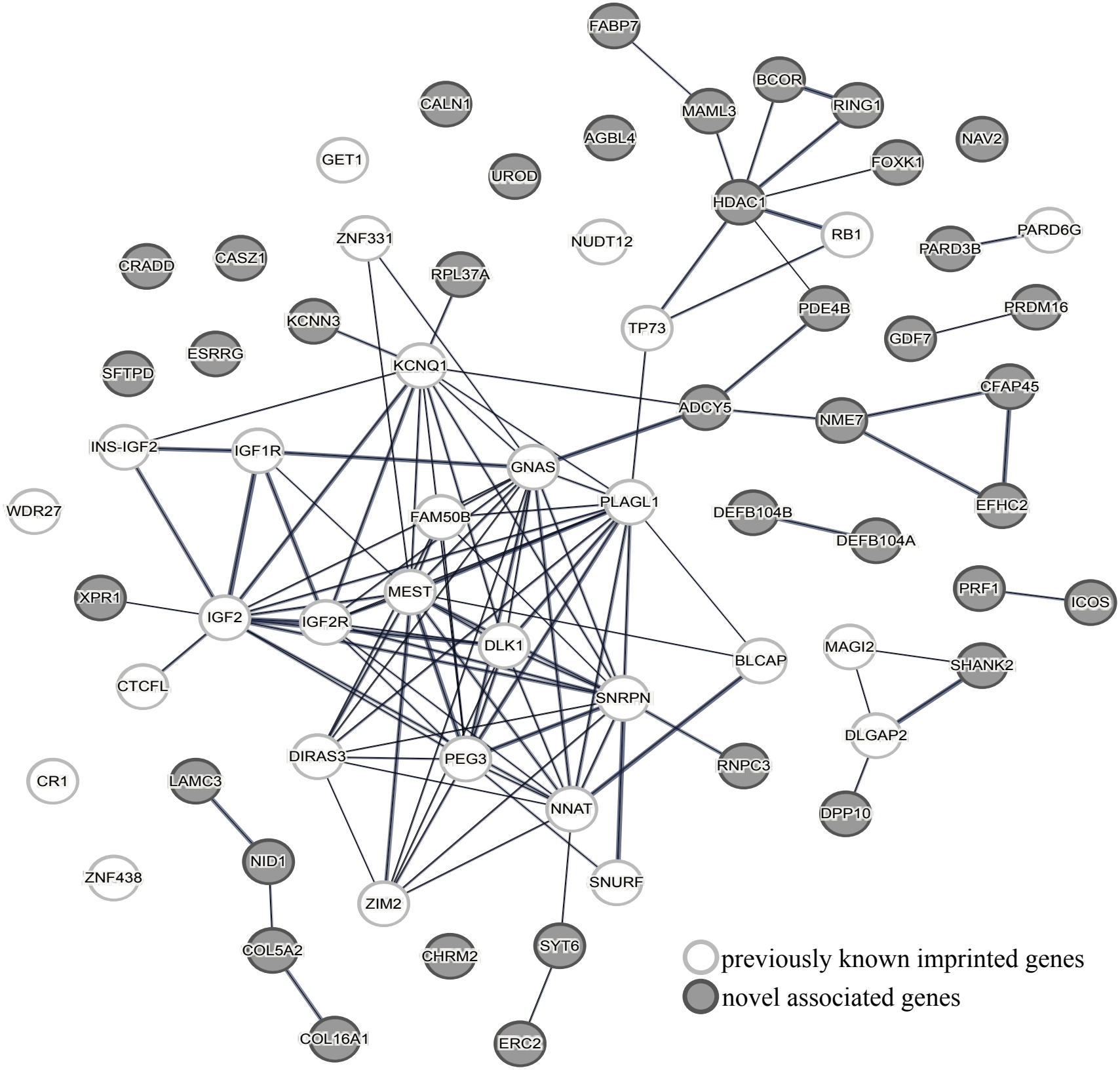
Protein–protein interaction network for genes associated with POE–dependent probes. The network is generated using STRING (22) and restricted to gene identifiers with interactions, association with the top six traits from phenotype enrichment analysis or previously mentioned in other studies. This results in a set of 68 genes associated with POE–dependent probes identified in our study. Novel associated genes are highlighted in gray and genes previously reported by Cuéllar-Partida *et al*. or Rosenski *et al*. are shown in white.

Interestingly, three probes (cg14772836, cg16051475 and cg16088894, shown in Supplementary Figure S5) are linked to *EFHC2*, a gene reported to escape X– inactivation (23). Escape is a precondition for detecting parent–of–origin effects on the X in our design: at genes subject to X–inactivation, random silencing of either X across cells averages out parent–of–origin differences in bulk methylation; whereas, at escape genes, both copies remain active and a parent–of–origin difference is measurable. *EFHC2* has also been implicated in symptom severity in Turner syndrome (24), a context in which X– linked parent–of–origin effects have been proposed.

We investigate whether the fine-mapped SNPs that we identify are associated with gene expression regulation by overlapping them with several large eQTL and pQTL databases: (i) OneK1K single–cell eQTL atlas (25); (ii) GTEx whole–blood eQTLs (26); (iii) UK Biobank plasma pQTLs (27); and (iv) POE–associated pQTLs and QTLs from Hofmeister *et al*. (13) using exact SNP identifiers or SNPs within linkage disequilibrium (LD *R*^2^ *≥*0.5 within 1 Mb). Of the 119 unique POE–dependent probes with fine–mapped SNPs, we overlap 47 (39.5%) with OneK1K, 47 (39.5%) with GTEx, and 11 (9.2%) with UK Biobank pQTLs. Conversely, of the 30 POE–QTLs and 7 POE– pQTLs reported by Hofmeister *et al*., we recover only 7 (23.3%) and 1 (14.3%) respectively.

Because most of these databases, like our own data, are derived from whole blood, many of the overlapping loci are, as expected, involved in immune cell regulation. We find matches with OneK1K eQTLs involving CD4 and CD8 T cells, NK cells, B cells and monocytes, associations to *CHMP1A, SPATA33, SPATA2L* and *CDK10* in GTEx, and recover immune related pQTLs for TNFRSF14 and ICAM3 from the UK Biobank.

Among the individual matches, several point to regulatory mechanisms. Of the 47 matches with GTEx, 26 are protein–coding genes, 17 lncRNAs and 4 related to pseudogenes. One SNP (rs1555405), which matched the UK Biobank plasma pQTLs, falls in a CTCF binding site, associated with imprinting control regulation (28). Furthermore, we find that rs67876192 acts as a coordinated eQTL for multiple *KRAB* zinc finger protein genes in both OneK1K and GTEx. These genes protect imprinted regions from demethylation (29), pointing towards a potential regulatory mechanism at these loci.

SNP rs12881545 is associated with three POE—dependent probes in our study (cg18279536 annotated to *DLK1*, and cg18538961 and cg06150437, both annotated to *MEG8* ; all three show maternal imprinting, see Supplementary Figure S7). This SNP is found in all four QTL datasets: as a cis—eQTL for *MEG3* in non– classical monocytes in OneK1K, as an eQTL for *BEGAIN* (a neighboring gene in the *MEG3* imprinting cluster), in the GTEx whole–blood data, as cis–pQTL for *DLK1* in the UK Biobank, and as a primary POE–pQTL for *DLK1* in Hofmeister *et al*..

### Indirect parental genetic effects at imprinted genes

Within the probes modeled with the *CMFI* framework, we identify 221 probes with significant indirect maternal genetic effects (*V*_*M*_ *>* 5%) and 489 with indirect paternal genetic effects (*V*_*F*_ *>* 5%). We refer to these as maternal indirect and paternal indirect dependent probes. A list of maternal and paternal dependent probes is given in Supplementary Dataset S4 and a comparison of the variances with significant effects is shown in Supplementary Figure S8.

Protein–protein interaction networks (Supplementary Figures S9 and S10, gene annotation available in Supplementary Dataset S3) show key genes involved in biologically coherent pathways such as aging, embryonic development, neurodevelopment, metabolism, epigenetic regulation, meiotic recombination and cellular stress responses. Since probes showing indirect parental genetic effects are obtained after the initial filtering using significant imprinting variance with the *CI* model, this enrichment may partly reflect overlap with imprinted genes rather than generalized indirect effects. A more extensive discussion of these results is given in the Supplementary Text.

Comparison of loci associated with indirect parental genetic effects and POEs resulted in three probes with all three significant effects (*MFI* ), 8 indirect maternal and POE–dependent probes and 16 indirect paternal and POE–dependent probes, suggesting overlapping but distinct biological processes (Supplementary Figure S11). Genomic imprinting relies on iDMRs, whereas indirect maternal genetic effects arise mainly through the intrauterine environment (30), indirect paternal effects through sperm and seminal plasma modifications (31–33) and both indirect parental genetic effects through the postnatal environment (34). Despite these mechanistic differences, loci associated with both indirect parental and imprinting effects indicate that both processes can interact and converge on common biological pathways.

In particular, genes inferred to be influenced by both POEs and indirect parental genetic effects were functionally enriched for terms related to aging (*P* = 3.49 *×*10^*−*3^), smoking (*P* = 7.21 *×*10^*−*3^) and gestational diabetes (*P* = 6.18*×* 10^*−*4^). This overlap is likely to be biologically relevant, as parental age and health both influence gamete quality (10, 31), and are associated with widespread epigenetic changes (8), including methylation modifications that contribute to the establishment and maintenance of genomic imprinting (35, 36). Moreover, imprinted genes play a central role in embryonic growth, placental function and metabolic regulation throughout life. These findings highlight the complexity of the mechanisms contributing to offspring DNA methylation variation and reinforce the importance of jointly modeling POEs and indirect parental genetic effects. Separating these sources of variation reduces the potential for confounding and provides a clearer framework for understanding the developmental origins of epigenetic variation.

## Discussion

Our results indicate that indirect parental genetic effects are an important source of apparent POEs in offspring methylation. By jointly modeling offspring and parental genotypes, we were able to separate the transmitted genetic component from indirect parental genetic effects and identify a substantially more restricted set of methylation sites with POEs. We further identified SNPs associated with those sites and compared the resulting signals with previous studies, recovering known regions, while also identifying putative novel signals. These loci included genes and downstream associations with gene expression, protein abundance, and phenotypes, providing additional context for the potential biological relevance of our POE–dependent methylation signals. At the same time, our analysis identified methylation sites that are potentially associated with indirect maternal and paternal genetic effects, highlighting an additional source of familial influence that may contribute to methylation differences attributed to POEs in previous studies. The reduction in POE–associated methylation sites after conditioning on parental genotypes suggests that estimates from designs lacking parental genotype information may overstate the extent of POEs in methylation.

There are several important limitations. Our study relies on both the methylation probe and the associated SNP having sufficiently strong and detectable signals. While our study is the largest available, sample sizes remain limited because genetic markers and methylation data are required for both parents and children. X chromosome analyses are restricted to the 1,182 trios with a daughter and are correspondingly less powered, so the 17 X–linked probes are likely a conservative subset. In addition, imprinting pattern classification depends on the estimated coefficients and variances of the associated SNPs, which restricts classification to probes with confidently associated variants and reduces the number of loci for which specific imprinting patterns can be inferred. Although including parental genomes in a parent–offspring trio design provides clear advantages for disentangling parental and imprinting effects, it also substantially restricts statistical power compared with standard population based studies. Furthermore, all samples are derived from a single cohort with relatively homogeneous ancestry and shared environmental back-grounds, which limits the generalization of our findings to other populations. Additionally, as no family cohort of comparable size has trio genotypes with offspring methylation, internal replication was the only available option. Finally, methylation is measured in whole blood from adult participants and thus our estimates represent the average pattern of imprinting across immune cell types in blood. As imprinting effects are often cell–type, tissue–, and development–stage–specific, relevant POEs present in other tissues or earlier stages of development may remain undetected and the patterns presented here may not hold across all instances. Despite these limitations, we have highlighted the importance of within–family designs for the identification of parent–of–origin effects: conditioning on trio genotypes gives POE estimates substantial protection against confounders such as cell composition, batch, ancestry, probe-level SNP artifacts and indirect genetic effects. These affect population-based methylation studies, and protecting against them is why this design is worth its cost in power.

We conclude that phenotypic variation is shaped by direct genetic, indirect parental and imprinting effects acting jointly. Accurately disentangling these contributions is essential for understanding how parental inheritance shapes the human epigenome. Studies of parent–of–origin methylation that do not condition on parental genotypes estimate a conflated upper bound rather than a quantity attributable to parental origin, a difference which may be large: in our data, nine in ten apparent signals do not remain. We do not assign these probes to a single cause; some probes will have indirect parental genetic effects that fall below our detection threshold, while others will reflect the reduced identifiability of a four–component model at this sample size. The practical consequence is the same in either case: existing catalogs of imprinted differentially methylated regions, and the downstream inferences built on them, warrant re–examination. The loci reported here should be read as a conservative, high–confidence set rather than an exhaustive one.

## Materials and Methods

### Generation Scotland

Generation Scotland (GS) is the largest within– family cohort where whole blood methylation, SNP genotypes and electronic health records are all available for 18,843 participants (37). The study baseline took place between 2006 and 2011 and included detailed cognitive, physical, and health questionnaires, along with sample donation for genetic and biomarker data. Ethical approval for the GS cohort was received from the NHS Tayside Committee on Medical Research Ethics (REC Reference Number: 05/S1401/89) and Research Tissue Bank status was granted by the East of Scotland Research Ethics Service (REC Reference Number: 25/ES/0013). Participants provided written informed consent.

Methylation was observed using the Infinium MethylationEPIC Bead-Chip array, comprising 850,055 methylation probes. The resulting data are adjusted for age, sex, ever smoked, packs per year and batch, standardized, and transformed using a rank–based inverse normal transformation to improve normality. To analyze the X chromosome, we restricted the samples to daughters (1,182 trios) and to non–pseudoautosomal regions (18,706 probes).

Genotyping array data are available from the Illumina Human Omni Express Exome8V.1–2 A and Human Omni Express Exome–8V.1 A arrays. We calculated the coefficient of relatedness among individuals with the software KING (38). Parents are identified as individuals of opposite biological sex who: (i) are unrelated (relatedness *<* 0.0012), (ii) are within 10 years of difference in age, (iii) both showed approximately 0.5 relatedness to the same individual, and (iv) are both at least 15 years older than the individual whom they share half their DNA with. The latter two criteria automatically selected the children. We then selected families for which methylation observations are available for the children, resulting in 2,438 parent-offspring trios. We additionally identified 3,854 parent– offspring duos and 4,856 sibling pairs from 13,564 individuals, whose relationship we also confirmed with the available family information.

We phased the array data using *SHAPEIT5 version 5*.*1*.*1* (39) using the *related individuals* option to leverage pedigree information. We then imputed the phased data with *IMPUTE5 version 1*.*1*.*5* (40) using the Haplotype Reference Consortium panel (HRC r1.1, GRCh37) as reference. For the X chromosome, phasing was performed using the options *pedigree* and *haploid*, and imputation was conducted separately for each sex. We removed samples with a call rate of *<* 98%, and variants with an INFO score *<* 0.9, minor allele frequency *<* 1%, and Hardy Weinburg equilibrium of *P <* 1 *×*10^*−*6^. After these quality measures, we obtained 11,358,693 imputed autosomal and 16,104 X–chromosome SNPs. We further restricted the number of SNPs to SNPs with minor allele frequency *>* 5%, *r*^2^ *≤* 0.8 and Mendelian error rate *<* 1%. Additionally, for the X chromosome, we excluded SNPs that showed no variation among fathers. This gives 931,607 autosomal and 5,371 X–chromosome SNPs.

### Study design

The methylation observations cannot be phased to maternally and paternally inherited chromosomes. Thus, to identify differentially methylated regions between maternal and paternal chromosomes, we modelled offspring methylation levels for each probe using a multivariable linear regression framework. The methylation values for a given probe, ***Y***, of *n* children were modeled as a linear function of the design matrix, ***X***, of size *n ×* 4*p*,

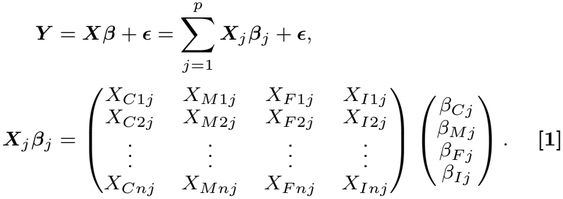

***X*** was composed of the child’s genotype *C*, the mother’s genotype *M*, the father’s genotype *F* and the parent–of–origin indicator *I*, with *j* = 1, …, *p* for all *p* SNPs present on the chromosome that the methylation probe was observed on, which avoids an initial arbitrary definition of the base–pair distance for ‘cis’ regions. ***ϵ*** represents the residual error. The effect of each SNP had four components, ***β***_*j*_ = (*β*_*Cj*_, *β*_*Mj*_, *β*_*F j*_, *β*_*Ij*_ )^*T*^, representing the direct genetic effects, *β*_*C*_, that children’s genotypes have on their own phenotype; the indirect parental genetic effects, *β*_*M*_ and *β*_*F*_, that parents’ genotypes have on their children; and parent–of–origin or imprinting effects (POEs), *β*_*I*_, that are dependent on the parental origin of the allele. Parental and child genotypes are coded 0 for homozygous reference allele, 1 for heterozygous, and 2 for homozygous alternative allele. The parent–of– origin indicator is 0 for homozygotes, *−*1 for heterozygotes inheriting the alternative allele from the father, and +1 for those inheriting it from the mother.

As parents and children share DNA, estimating the direct effects of a child’s DNA requires conditioning on the variants in the parents. Since parents are the source of genetic variation in their children, jointly modeling parental and offspring genotypes allows us to account for indirect genetic effects arising from correlations between parental and offspring genotypes. In addition, because the potential genotype of the child is strictly limited by the combinations of alleles available in the parents, conditioning on the variants in the parents is required to account for the correlation of the parental genotypes and the indicator for the parent–of–origin of each SNP inherited by reciprocal heterozygotes. This conditioning is necessary to account for the correlation between parental genotypes, offspring genotype, and parent–of–origin indicators in this design. Because we modelled the four genetic components of all SNPs together, the number of predictors 4*p* exceeds the number of observations *n*, resulting in an under–determined high–dimensional regression setting. Thus, we first evaluated three potential penalized linear regression approaches using simulated data: JODIE (18), which has recently been proposed for estimation of direct and indirect effects as well as POEs; SuSiE (41), a commonly employed variational inference fine–mapping approach; and the LASSO model (42), the most commonly applied penalized regression model, with the goal of simply selecting the most appropriate approach for the GS trio data. The models and their performance are discussed more extensively in the Supplementary Text.

### Simulation study

We simulated phenotypes based on real GS genotype data from chromosome 22 (16,466 SNPs) of 2,438 parent–offspring trios. For each simulation replicate, five SNPs with minor allele frequency between 0.3 and 0.5 and pairwise *R*^2^ *<* 0.2 were randomly selected and assigned non–zero effects. Their effect sizes were drawn from a multivariate normal distribution with mean 0 and covariance:

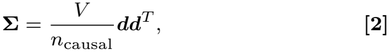

where the variance *V* was set to 0.2 (unless otherwise indicated), and is equally distributed across *n*_causal_ = 5 causal SNPs. The vector ***d*** reflects various imprinting patterns, which are summarized in Table 1. Phenotypes were generated as:

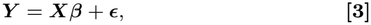

**Table 1.** Simulation scenarios. Simulated patterns for each scenario and the signs of each genetic effect (*d*) with order (C, M, F, I).

| **Pattern** | $\boldsymbol{d}$ |
|---|---|
| Positive paternal imprinting | $(+1, 0, 0, +1)^T$ |
| Negative paternal imprinting | $(-1, 0, 0, -1)^T$ |
| Positive maternal imprinting | $(+1, 0, 0, -1)^T$ |
| Negative maternal imprinting | $(-1, 0, 0, +1)^T$ |
| Positive complex imprinting | $(0, 0, 0, +1)^T$ |
| Negative complex imprinting | $(0, 0, 0, -1)^T$ |
| Direct and indirect maternal | $(+1, +1, 0, 0)^T$ |
| Direct and indirect paternal | $(+1, 0, +1, 0)^T$ |
| Indirect maternal effect | $(0, +1, 0, 0)^T$ |
| Indirect paternal effect | $(0, 0, +1, 0)^T$ |
| Indirect maternal and paternal effects | $(0, +1, +1, 0)^T$ |
| Direct genetic effect | $(+1, 0, 0, 0)^T$ |

where ***X*** is the standardized genotype data and ***ϵ*** is a random error term drawn from a normal distribution with mean 0 and variance *σ*^2^ = 1 *−*Var(***Xβ***).

To evaluate the ability of JODIE, SuSiE, and LASSO to correctly identify causal SNPs and their corresponding effects while minimizing false positive imprinting signals, we generated 400 datasets in total, simulating different scenarios of imprinting as well as direct and indirect parental genetic effects (given in the first 8 rows of Table 1).

For LASSO, we used 100 cross–validation steps to identify the optimal regularization parameter, then applied stability selection retaining coefficients present in at least 95 of 100 runs, with the average coefficient estimate used. For SuSiE and JODIE, coefficients with PIP *≥* 0.95 were set to zero and posterior mean estimates are used. JODIE additionally provides variance estimates for each of the four effect components,accounting for assortative mating (18). We applied a threshold of *>* 5% on the variances. These criteria ensures that probes are classified as having significant effects only when the corresponding effects are distinguishable from zero and from competing associations (Supplementary Figure S12). Performance was evaluated using: (i) Pearson’s *R*^2^ between true (***β***) and estimated 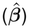 coefficients; (ii) true positive rate (TPR) calculated as the ratio of identified true causal variants to the total number of simulated causal variants; and (iii) false discovery rate (FDR) calculated as the proportion of false positives among all hypotheses declared to be significant. These three performance metrics were computed both per component and per SNP, to examine the ability of the models to recover the correct SNP and the correct simulated effect type. Results are given in Supplementary Figure S13. JODIE demonstrates the best performance, particularly in minimizing false discoveries.

To ensure that these conclusions remain valid across different values of *V*, we generated 20 datasets each for *V* = 0.1, 0.2, 0.3, 0.4, using only direct genetic effects (last row of Table 1), whilst ensuring that the simulated *V* stays constant across replicates. The TPR and FDR as a function of *V* are shown in Supplementary Figure S14. Based on these results, JODIE was selected for the analysis of real data.

Additionally, we wanted to demonstrate that indirect parental genetic effects can mimic true imprinting when parental genotypes and parent– of–origin indicators are not modeled jointly. For this, we simulated 50 datasets each, where only indirect maternal genetic effects, only indirect paternal genetic effects, and both maternal and paternal effects exist. We analyzed the data using JODIE: first, restricted only to child’s genotypes and parent–of–origin indicator (*CI* model), and then including also maternal and paternal genotypes (*CMFI* model). The comparison of the two models shows that imprinting effects and indirect parental genetic effects need to be modeled jointly to ensure an unbiased estimation of these effects (Supplementary Figure S1).

### Identification of genomic regions with differential methylation between maternal and paternal chromosomes

To reduce the computational runtime of the iterative Bayesian sampler, we implemented a two–step screening framework using JODIE: first, a reduced model using only the child’s genotype and parent–of–origin indicators (*CI* model) was run for 1000 iterations, independently for each of the 850,055 methylation probes using all SNPs on the corresponding chromosome. The number of SNPs and probes per chromosome is shown in Supplementary Figure S15.

From the 787,909 of 850,055 models that converged, putative POE– dependent probes were identified by having a substantial amount of variance attributed to POEs (*V*_*I*_ *>* 5%). Because the rank based inverse normal transformation alters the scale of the methylation phenotypes, all estimated variances were normalized by the maximum observed phenotype value. This normalization value was also used to filter out probes with high uncertainty in their estimated variance values. We identified 5,435 putative POE–dependent probes during step 1.

In a second step, we re–analyzed the 5,435 putative POE–dependent probes for 3000 iterations using the full *CMFI* model, where child, maternal and paternal genotypes are modeled jointly alongside the parent–of–origin indicators.

Thus, we tested whether the variance attributed to POEs remains after conditioning on maternal and paternal indirect genetic effects. Of the 5,435 probes, 59 did not converge; for these, we found no association with any factor besides chromosome position, with X-chromosome probes accounting for 40.7% of the total non–converged probes, suggesting that our results are not biased but rather restricted by sample size and reduced power. We applied an additional filter on the uncertainty of the estimated variance components, excluding a further 121 probes. Of the remaining 5,255 probes, 4,974 (94.6% of remaining probes and 91.5% of the 5,435 putative POE–dependent probes) fell below the 5% variance threshold, and 281 remained as POE–dependent probes, of which 17 are found on the X chromosome (Figure 2). Similarly, we also identified probes with indirect parental genetic effects conditional on POEs by applying a 5% threshold on the variances attributable to indirect maternal and paternal effects. We identified 221 putative probes (including 5 on the X chromosome) that are dependent on indirect maternal genetic effects, and 489 that are dependent on indirect paternal genetic effects. The larger number of indirect paternal probes compared to indirect maternal probes is unlikely to reflect a true biological asymmetry. Rather this is a consequence of the initial filtering for *V*_*I*_ *>* 5% applied to the *CI* model, which presents a higher ratio of probes exhibiting methylation patterns consistent with maternal imprinting (paternal expression), subsequently identifying more indirect paternal genetic effects when fitting the *CMFI* model (Supplementary Figure S16).

This two–step framework substantially reduced computational cost, while enabling direct comparison of imprinting estimates with and without parental effects. To further validate the estimated coefficients, we compared the results from the *CMFI* model with those obtained using SuSiE applied to a reduced set of cis–SNPs (*<* 5 Mb). The results are presented in Supplementary Figure S17, which shows how the estimates are strongly correlated, although some effects detected by JODIE are not found by SuSiE, potentially reflecting differences in statistical power between methods, shown in Supplementary Figure S13.

### Fine–mapping SNPs within imprinted methylated regions

We performed fine–mapping to identify candidate causal variants within genomic regions. From the *CMFI* model, we select a SNP *j* SNP as being associated with POE–dependent probes when the posterior inclusion probability was PIP *≥* 0.95, the variance in any of the *k* components was *>* 5% and the estimated SNP effect in that component was 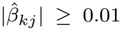 . This is conservative, but justified under the assumption that individual SNP components are unlikely to exert substantial phenotypic effects when the total variance explained by that component is negligible and we wished to focus on the leading associated regions.

The genomic distance between each probe and its associated SNPs is small, with only a handful of variants located more than 1 Mb away (Supplementary Figure S18), justifying a 2 Mb fine–mapping window. Thus, for each identified POE–dependent probe and each associated SNP, all imputed SNPs within a 2 Mb window, with minor allele frequency *>* 5%, and Mendelian error rate *<* 1%, but without applying the strict *R*^2^ *≤* 0.8 filter, were re–analyzed using the *CMFI* model.

To account for the high correlation structure among nearby SNPs, we grouped the posterior distribution into credible sets, defined as sets of SNPs with pairwise *R*^2^ *≥* 0.5. The posterior inclusion probability of each credible set was computed as the proportion of posterior iterations in which at least one SNP within the credible set was included in the model. Credible sets with a PIP *≥* 0.95 were retained as significant and SNPs included in at least 5% of the total number of iterations of the credible set were considered fine–mapped.

### Classification of imprinting patterns

The parent–of–origin indicators provide information about which parent has contributed the alternative allele in case of heterozygous children (*X*_*I*_ = +1 for maternally inherited, and *X*_*I*_ = *−*1 for paternally inherited alleles), but not which parental allele is being expressed. To be able to classify imprinting patterns, we therefore also need the estimated direct effects to be able to distinguish the cases where the alternative allele results in higher (*β*_*C*_ *>* 0) and lower methylation levels (*β*_*C*_ *<* 0) compared to the reference allele.

For example, consider a locus with *β*_*C*_ *>* 0. A heterozygote who inherited the alternative allele from the mother has *X*_*C*_ = 1 and *X*_*I*_ = +1, and therefore an expected methylation level of *β*_*C*_ + *β*_*I*_ ; the reciprocal heterozygote has *X*_*C*_ = *−*1 and *X*_*I*_ = *−*1, and an expected level of *β*_*C*_ *β*_*I*_ . When *β*_*I*_ *>* 0 and of comparable magnitude to *β*_*C*_, the paternally inherited alternative allele therefore behaves as if it were the reference allele, while the maternally inherited copy carries the full effect: the cis effect of the variant on methylation is expressed only on the maternal copy. When *β*_*I*_ *<* 0, the pattern is reversed and the effect is expressed only on the paternal copy.

We interpreted the unresponsive copy as the imprinted one. A copy that carries a constitutive parent–of–origin methylation mark has little dynamic range remaining for a cis variant to act on, so the effect of that variant is detectable mainly on the copy that is not marked. Under this assumption, *β*_*C*_ and *β*_*I*_ sharing a sign identifies the paternally inherited copy as the silenced one (paternal imprinting, with expression from the maternal allele), and opposite signs identify the maternal copy as silenced (maternal imprinting, expression from the paternal allele). The classification itself rests only on methylation; mapping the silenced copy onto an expressed allele is the single additional assumption.

When a significant imprinting effect is detected while the direct effect *β*_*C*_ is indistinguishable from zero, the observed pattern can be attributed to dominance mechanisms, such as over–dominance, under–dominance, or bipolar dominance patterns, which we refer to as complex imprinting patterns (Figure 3). The possible imprinting patterns and their classification based on the direction of direct and imprinting effects are summarized in Table 2. In contrast, non–imprinting patterns are characterized by assessing whether the interval estimates for direct *β*_*C*_ and indirect parental genetic effects *β*_*M*_ and *β*_*F*_ are different from zero.

**Table 2.**
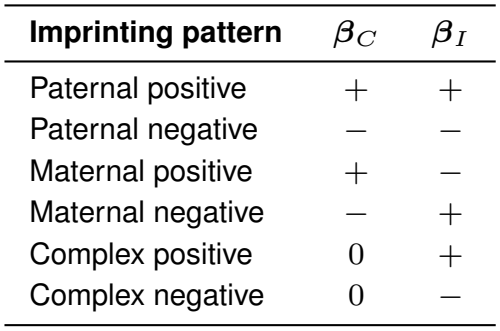
Classification of imprinting patterns. Imprinting patterns are inferred from the signs of the direct genetic (*β*_*C*_ **) and imprinting (***β*_*I*_ **) effects. Positive and negative signs indicate the direction of the estimated effect, whereas** 0 **denotes a direct effect indistinguishable from zero**.

| **Imprinting pattern** | **β**<sub>*C*</sub> | **β**<sub>*I*</sub> |
|---|---|---|
| Paternal positive | + | + |
| Paternal negative | − | − |
| Maternal positive | + | − |
| Maternal negative | − | + |
| Complex positive | 0 | + |
| Complex negative | 0 | − |

Only fine–mapped SNPs were used for the classification of imprinting patterns. For each iteration, where the fine–mapped SNP was included in the model, the imprinting pattern was identified according to Table 2. When a single SNP was included in several models (due to an overlap of fine–mapping windows), we used the estimates and iterations from all models. If the same imprinting pattern occurred in at least 75% of the iterations, the SNP was classified according to this pattern, otherwise the SNP was classified as undefined.

Average methylation values ***Y*** ^*′*^ for individual SNPs *j*, such as displayed in Figure 4, were calculated by residualizing the normalized and rank– inverse–transformed phenotype against the estimated effects of all other variants:

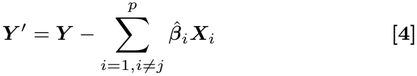

When adjusting for a specific component of SNP *j*, we also residualized the phenotype against all other components of *j*, so that only the contribution of the component remained.

### Gene annotation

To annotate methylation probes with significant effects, we used Illumina Human Omni Express Exome8V.1–2 A and Human Omni Express Exome–8V.1 A array annotation files and the Methylation Screening Array (MSA) hg38 gencode v41 (43). Associated fine– mapped SNPs were annotated using the Ensembl Variant Effect Predictor (VEP) (44) and used as supporting evidence for genes annotated from POE– dependent probes. We generated protein–protein interaction networks utilizing gene identifiers from both Illumina array annotation files and MSA and STRING (22) with confidence based edges with a minimum interaction confidence score of 0.4 and restricting displayed nodes to those with connections. Furthermore, we performed an enrichment annotation for those probes overlapping multiple effects using the EWAS catalogue (45) and EWAS Toolkit with EPIC/850K background (46).

### Replication and comparison to other studies

We compared the POE– dependent probes against those identified by previous studies including: parental allele specific methylation and known imprinted differentially methylated regions from Rosenski *et al*. (15); known imprinted regions and POE–dependent probes reported by Cuéllar-Partida *et al*. (17); and POE–influenced probes identified in a previous analysis in the Generation Scotland cohort by Zeng *et al*. (16).

Probes were replicated using both probe identifiers when available, and genomic location (distance *≤* 2 Mb), this threshold was selected for being conservative towards novel probes and for consistency between studies since it was also used by Cuéllar-Partida *et al*.. To assess differences in the estimated variance distributions between novel probes, overlapping probes, and probes reported only in these other studies, we performed Mann– Whitney U tests for each variance component and obtained corresponding *P− values* for pairwise comparisons.

Under imprinting, siblings and parent–offspring duos are expected to resemble each other more closely than predicted by additive genetic effects alone. We leveraged this expectation to replicate findings from the trio based analysis by extending the approach described in Zeng *et al*. This framework compares a set of alternative models with different relatedness matrices ***S***:

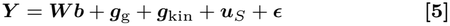

with ***W*** a matrix of covariates (age, sex) and ***u***_*S*_ representing a random effect given by POE:

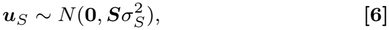

against a null model which includes only additive genetic 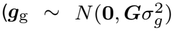, with ***G*** a relationship matrix calculated from the SNPs) and pedigree 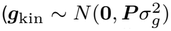, with ***P*** a relationship matrix calculated from the pedigree) random effects along with the fixed effects ***Wb***:

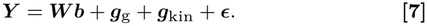

We used four relatedness matrices to define ***S***, each representing different family resemblance structures: siblings (***S***_*s*_), siblings and mother– offspring duos (***S***_sm_), siblings and father–offspring duos (***S***_sp_), and siblings with either parent–offspring duo (***S***_smp_). Model comparisons are performed using likelihood ratio tests (LRTs) calculated with *GREML* (47). To ensure robust model selection, we applied the following strategy: when the constrained and unconstrained analyses identify the same best-fitting model, that model was selected. When they disagree, the constrained model is preferred unless the unconstrained model shows a stronger fit (higher LRT). Probes for which an alternative model was selected as opposed to the null model were classified as POE–dependent and considered replicated as shown in Figure 5c.

### Data, Materials, and Software Availability

Access to the Generation Scotland data is available with appropriate permission from the Generation Scotland Access Committee. Access requests can be made through the researcher portal (https://gsaccess.igc.ed.ac.uk) and normally take up to six weeks for approval. Further information can be found on the Generation Scotland website: https://genscot.ed.ac.uk/for-researchers/access. Haplotype Reference Consortium Release 1.1 data (https://ega-archive.org/datasets/EGAD00001002729) are available by application to a Data Access Committee (DAC) of the Wellcome Trust Sanger Institute.

#### Code availability

1. Custom Bayesian model code: https://github.com/medical-genomics-group/JODIE.
2. Analysis pipeline and code: https://github.com/medical-genomics-group/disentangling_genomicImprinting_indirectEffects.

## ACKNOWLEDGMENTS

We thank and acknowledge the participants and investigators of the Generation Scotland Cohort study. Generation Scotland received core support from the Chief Scientist Office of the Scottish Government Health Directorates [CZD/16/6] and the Scottish Funding Council [HR03006]. Genotyping and methylation typing of the GS:SFHS samples was carried out by the Genetics Core Laboratory at the University of Edinburgh Clinical Research Facility, Edinburgh, Scotland and was funded by the Medical Research Council UK and the Wellcome Trust (Wellcome Trust Strategic Award “STratifying Resilience and Depression Longitudinally” (STRADL) Reference 104036/Z/14/Z). DNA methylation profiling was also funded in part by the Wellcome Trust Investigator Award (Reference 220857/Z/20/Z). GS is currently supported by the Wellcome Trust (216767/Z/19/Z).

Analysis of the Generation Scotland data and the summary statistics obtained from the other analyses was conducted at IST Austria and is supported by the Scientific Service Units (SSU) of IST Austria through resources provided by Scientific Computing (SciComp).

