## Supplementary Information for "Disentangling genomic imprinting from indirect parental genetic effects on the human methylome"

Fernanda Adame Galeano[a], Ilse Krätschmer[a], Anne Richmond[b], Caroline Hayward[b], Riccardo E. Marioni[b], Beatriz Vicoso[a], Matthew R. Robinson[a]
[a] Institute of Science and Technology Austria, Klosterneuburg, Austria, 3400
[b] Institute of Genetics and Cancer, University of Edinburgh, Edinburgh, UK, EH4 2XU

November 17, 2026

#### Supplementary Text

##### Simulation study results

We find that JODIE outperforms SuSiE and LASSO models across all settings (Figure 13). These findings remain the same when testing whether the methods correctly attributed detected signals to the appropriate biological component, by either: (i) assessing whether the correct SNP and effect type ($C$, $M$, $F$, or $I$) were recovered; or (ii) assessing whether the correct causal marker was identified, regardless of which component carried the effect. Although all approaches identify the correct SNP with high precision across these settings, LASSO and SuSiE occasionally misattribute effects within the same SNP, for example assigning the signal to the direct rather than the imprinting component as they are not multi–component models. Additionally, as the heritability differs across simulation settings, we explore the empirical power (TPR) and FDR in Figure 14, where we find that JODIE maintains a high TPR and low FDR across all heritability and simulated effect size magnitudes.

Uniquely among the approaches, JODIE is a grouped regression model that allows covariances between POE, direct, and indirect effects to be estimated chromosome–wide across SNPs, which enables the pattern of imprinting to be inferred. We show in Figure 2 that JODIE not only separates the variance attributable to each source of genetic influence, but also preserves the expected directional relationship between direct and imprinting effects, estimating the direction and magnitude of the covariance patterns accurately. Thus, we confirm that, for these data and under these parameter and simulation settings, JODIE fine–maps genetic effects accurately, whilst minimizing false positives, and correctly identifies imprinting patterns.

In an additional simulation study comprised of 150 datsets, we simulate three effect patterns where there are no POEs and: (i) only indirect maternal effects; (ii) only indirect paternal effects; and (iii) both indirect maternal and paternal effects. For each scenario, we fit a design matrix containing only direct and POE encodings for all SNPs (termed "$CI$ model") and a model where the design matrix contains all four encodings for all SNPs (termed "*CMFI* model"). In Figure 1, we show that parental genotype effects are allocated to the inferred direct and imprinting effect variances of the *CI* model, which results in false identification of differentially methylated regions between maternal and paternal chromosomes. We further demonstrate that parental genetic effects mimic the phenotypic patterns of imprinting expected for the average phenotypic values in Figure 4b, with a difference between reciprocal heterozygous observed despite there being no simulated POE in this scenario. However, across all settings once parental genotypes are jointly modeled, POEs are inferred to be zero (Figure 1). Taken together, our simulation results show that the effects of the different components can be disentangled within these data using JODIE, enhancing the accuracy of POE detection, and improving the interpretation of genetic associations.

#### Indirect parental genetic effects

By jointly fitting parental genotypes and parent–of–origin indicators using the *CMFI* model, we are able to identify probes that have significant variance due to indirect maternal and paternal effects ($V_M > 5\%$ or $V_F > 5\%$).

We find 221 and 489 probes with significant variance due to indirect maternal and paternal effects, respectively. As discussed in the main text, this is most likely a consequence of our initial filtering for $V_I \geq 5\%$ applied to the *CI* model, which distribution as seen in Supplementary Figure 16 shows more probes aligned with paternal expression. Similarly, because the enrichment analysis was conducted against a standard genome–wide background, we cannot rule out that some of the observed pathway enrichment reflects properties of imprinted genes, rather than independent indirect parental effects.

47 probes show significant variances due to both indirect maternal and paternal effects, but not POEs, as shown in Supplementary Figure 11. According to the EWAS catalog [1] and EWAS Toolkit [2], several of these probes are associated with aging, multiple syndromes, prenatal and postnatal exposures, such as maternal alcohol consumption, arsenic, assisted reproductive technologies, childhood adverse experiences, gestational age and gestational diabetes. A notable example is cg11666770 associated with smoking, differential methylation for Down syndrome newborns, diet and physical activity intervention for gestational diabetes, gestational age, differential methylation based on sex of infants and aging [3, 4, 5, 6, 7, 8]. Further three probes, namely cg18939097, cg18869304 and cg13706404, exhibit large variances attributed to both indirect parental effects and POEs. cg13706404 is associated with differential methylation in Down syndrome [6].

Additionally to these three probes, 8 POE–dependent probes also have $V_M > 5\%$, out of which 5 are situated on the X chromosome, an interesting pattern potentially linked to X–inactivation and escape; however, considering the small overlap this warrants further investigation. One of these overlapping probes is cg21028156 associated with differential methylation on umbilical cord of infants born from women with gestational diabetes [9].

Another 16 POE–dependent probes have $V_F > 5\%$, one of which is cg09395443 associated with gestational age and aging during childhood [5, 8]. cg09395443 is linked to *FOXK1*, a regulator of metabolism, autophagy, differentiation and known to demethylate in sperm with paternal age [10].

Associations with probes that have both variances due to POEs and one of the indirect parental effects include aging, metabolic phenotypes, smoking, alcohol consumption, infertility, gestational age, childhood associations with BMI and insulin sensitivity and immune phenotypes.

Finding probes with variances due to indirect maternal and paternal effects enriched for aging (p=$2.03 \times 10^{-13}$) suggests that parental genotypes influence the epigenetic aging trajectory of the offspring. The role of epigenetics and in specific methylation in aging has been well characterized [11]. Although transgenerational programming of longevity has been documented in animal models, characterizing such effects in humans remains a challenge [12].

Fine–mapping of the maternal and paternal dependent probes did not identify significant SNPs. This is likely due to the limited sample size along with the substantially smaller variance explained by indirect parental effects compared to imprinting effects (Supplementary Figure 8). This is biologically expected as imprinting operates through germline established iDMRs resulting in monoallelic expression, producing large effects, whereas indirect parental effects act through various physiological and environmental mechanisms from parent to offspring, caused by polygenic signals and resulting in smaller contributions to the variance.

Gene annotation identifies 167 unique gene identifiers associated with maternal dependent probes and 395 associated with paternal dependent probes. We generate protein–protein interaction networks from these genes (restricted to only genes with interactions) using STRING [13].

The network found for genes linked to maternal dependent probes, shown in Supplementary Figure 9, includes 50 genes in multiple small groups. The largest group comprises components of Sonic Hedgehog and insulin signaling pathways, such as *GLI2*, *SMO*, *LHX4*, *FOXO1* and *YWHAE*. These genes regulate embryonic patterning, organogenesis and metabolic signaling, and have previously been shown to respond to maternal environmental conditions during pregnancy, such as nutrition and diabetes [14, 15, 16]. Within that group we also find focal adhesion and cytoskeletal regulators (*PTK2*, *TNS1* and *ARHGEF1*) connected with each other, which control cell migration, extracellular matrix interactions and cytoskeletal remodeling, processes that are essential for placental development and labor [17, 18, 19, 20]. A second group of genes, which includes the voltage–gated calcium channels *CACNA1D* and *CACNA2D1*, the potassium channel *KCND2*, and the synaptic vesicle regulator *SYT1*, is enriched for genes involved in neuronal excitability and synaptic transmission. Together these genes regulate calcium–dependent neurotransmission, a pathway consistently implicated in maternal immune activation and prenatal inflammatory exposures associated with altered offspring neurodevelopment [21, 22]. The network also includes the X–linked calcium–binding protein *EFHC2*, which is not only associated with maternal indirect effects, but also POEs in our analysis. Finally, we also find *PRDM9* within the network, the primary determinant of meiotic recombination hotspots [23].

The network for genes linked to paternal dependent probes is shown in Supplementary Figure 10. It includes 174 genes and is centered on *AKT1*, a master regulator of cell survival, proliferation and metabolism that is also required for normal spermatogenesis and sperm quality [24]. We find several developmental regulators also identified in the maternal network, such as *GLI2*, *SMO* and *FOXO1*, which in experiments with mice, have been implicated in spermatogenesis and paternal dietary effects on offspring insulin sensitivity [25, 26, 27, 28]. Moreover, we also find the voltage–gated calcium channels *CACNA1D* and *CACNA2D1*, the potassium channel *KCND2*, and the synaptic vesicle regulator *SYT1* that we already identified in the network of genes linked to maternal dependent probes. Interestingly, *SYT1* has been previously associated with indirect paternal effects in experiments with mice, where spatial paternal training improves offspring's cognitive performance through changes in sperm [29, 30].

There are several genes from the HOXA family (*HOXA3*, *HOXA4*, *HOXA5*, *HOXA9* and *HOXA11*) present in the network for genes linked to paternal dependent probes, consistent with previous studies demonstrating paternal epigenetic regulation of embryonic body–axis specification genes [31, 32]. Another cluster comprises heat–shock proteins (*HSPA8*, *HSPD1* and *HSPE1*), which are essential for mitochondrial protein homeostasis, sperm maturation and fertility, and have been proposed as mediators of environmentally induced paternal epigenetic inheritance [33, 34, 35]. The network also includes key chromatin and germline regulators, such as *PRDM1*, *NSD1* and *NSD2*, all involved in epigenetic programming during development and germ–cell specification [36, 37], along with meiotic regulators, like *HFM1* [38]. Moreover, *AGO2* has previously been linked to POEs and is a key regulator of degradation of maternal mRNA and sperm quality [39, 40, 41]. Finally, we also find *MAGI2* within this network, which we already previously linked to the POE–dependent probes found in our study. It has been shown that *MAGI2* changes offspring methylation with paternal overweight [42].

information)

### 1 Supplementary figures

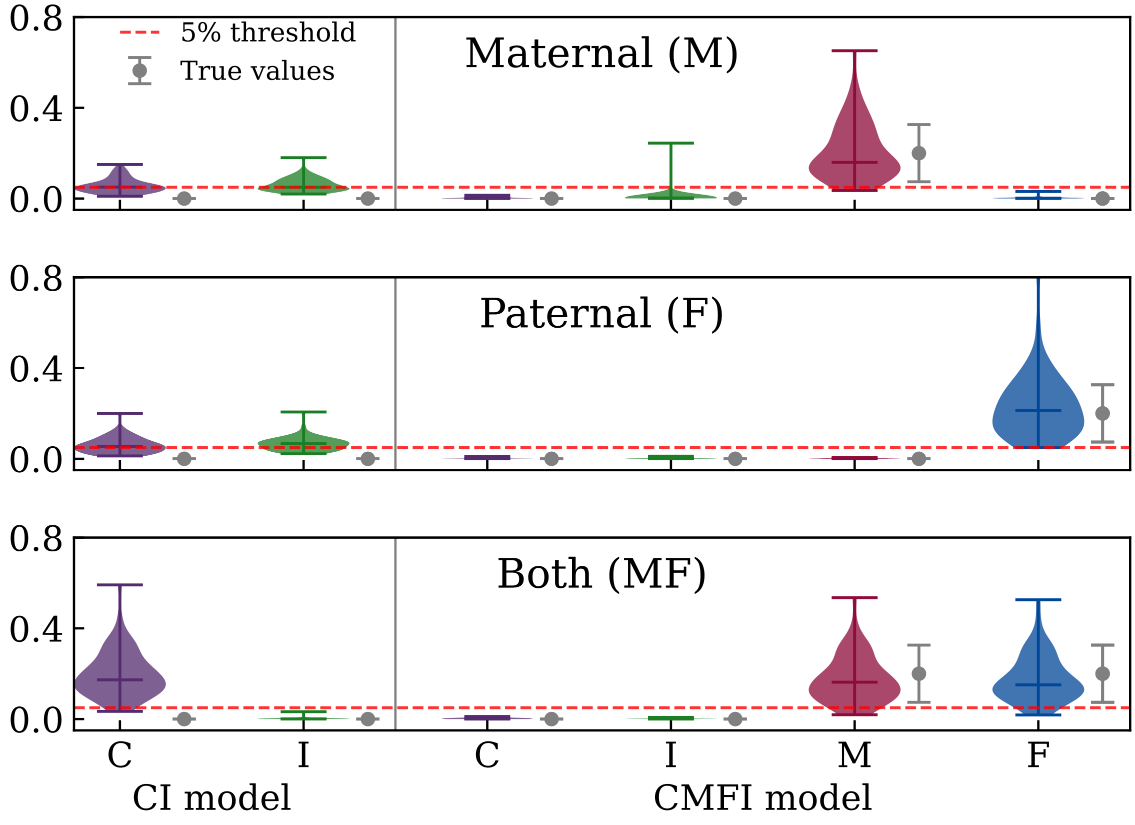


**Figure 1: Violin plot of variances estimated under the *CI* (left) and the *CMFI* model (right) using simulated data.** 50 replicates are generated for each simulation scenario: only indirect maternal effects (top), only indirect paternal effects (middle), and both maternal and paternal effects (bottom). When indirect parental effects are not modeled, their explained variance gets allocated to direct genetic and/or imprinting effects.

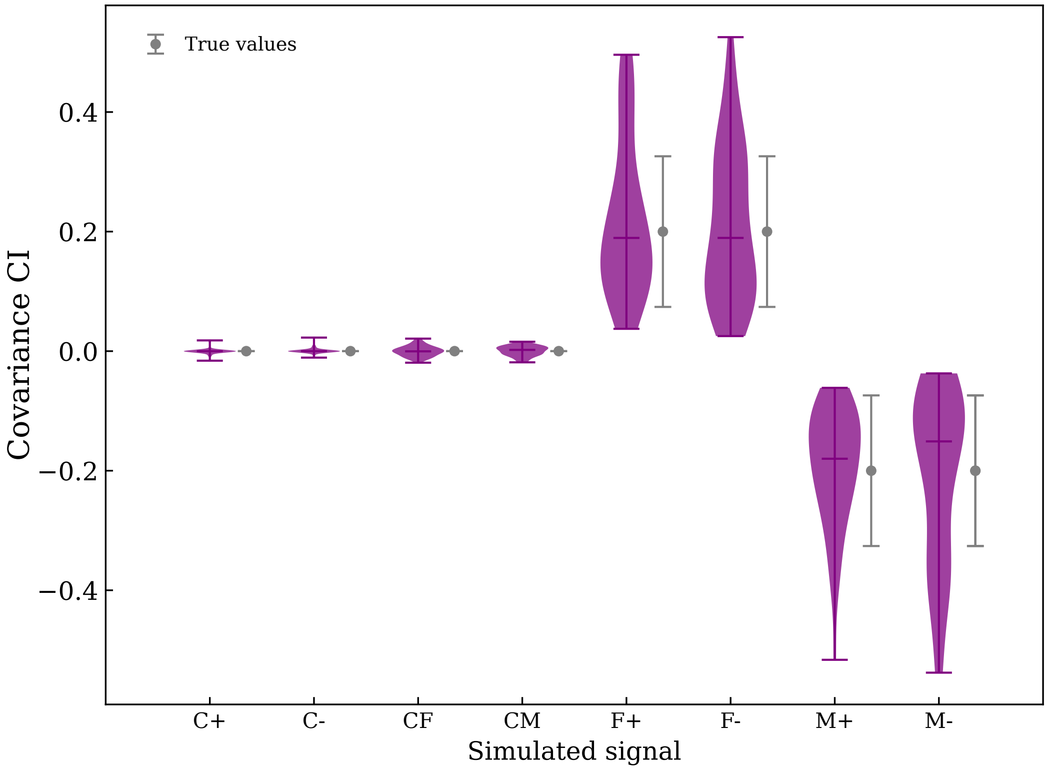


**Figure 2: Violin plot of estimated covariances between direct effects and POEs using the *CMFI* model for different simulated imprinting patterns.** 50 replicates are simulated for each of the eight patterns, shown in Table 2. In purple the inferred distribution of covariances between direct and imprinting components given by JODIE, in grey the expected values based on the simulated variance and standard deviation of the sampling distribution. First two distributions simulate complex imprinting patterns, CF and CM direct plus indirect paternal and maternal effects respectively and the last 4 include parental imprinting effects.

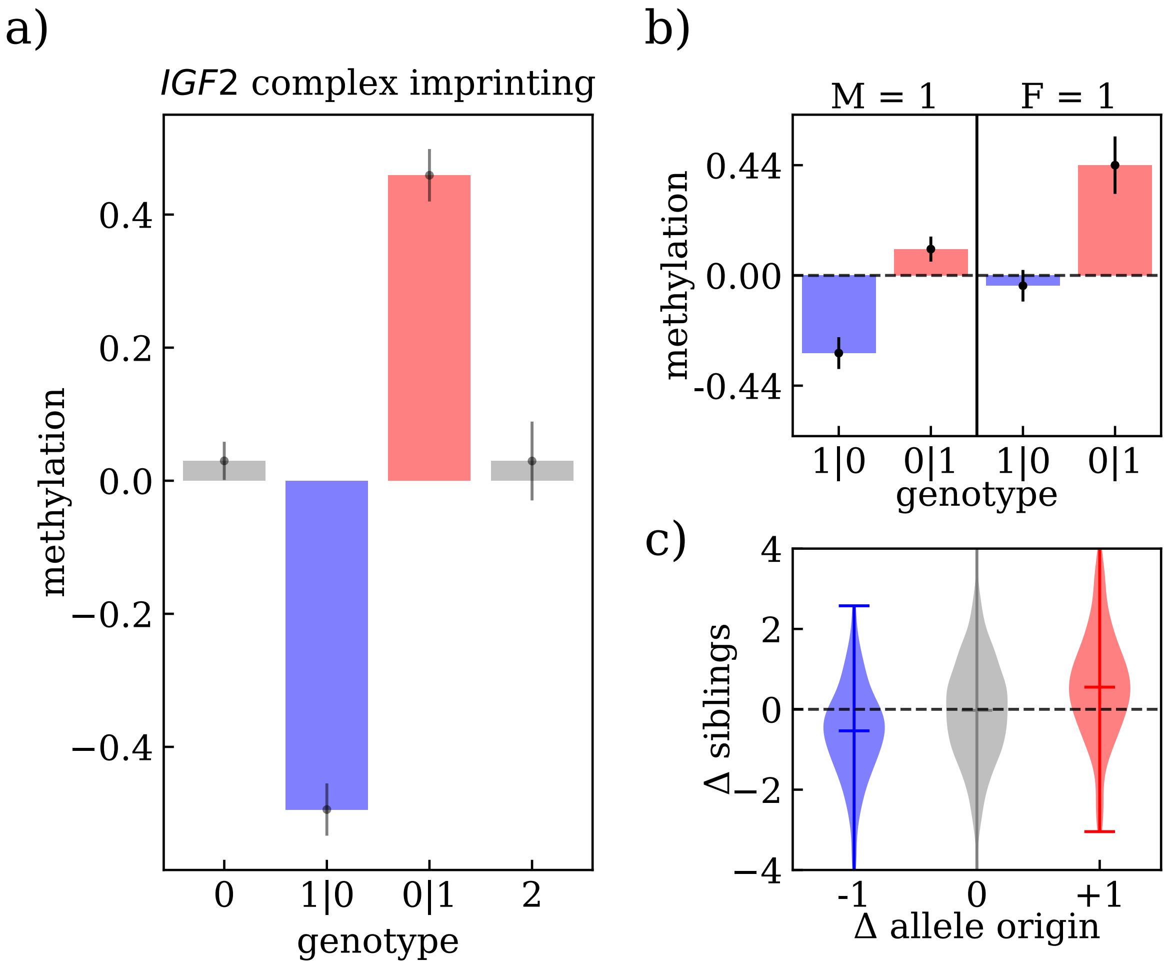


**Figure 3: Methylation patterns for rs7481173 acting on probe cg02425416, associated with *IGF2/INS-IGF2*.** (a) Average methylation of all children adjusted for the selected SNP and stratified by genotype, where differences between reciprocal heterozygotes (shown in blue and red) are typically interpreted as evidence of imprinting. (b) Average methylation of reciprocal heterozygotes from heterozygous parents adjusted for the indirect maternal $M$ or paternal $F$ component of the SNP, where similar measurements for both reciprocal heterozygotes from either homozygous mothers or fathers point towards indirect parental effects rather than imprinting. (c) Violin plots of differences in methylation of sibling pairs adjusted for imprinting effect on the SNP, stratified according to parental origin of the alternative allele, where a slope between different sibling pairs indicates an imprinting effect. Error bars in (a) and (b) correspond to the standard deviation.

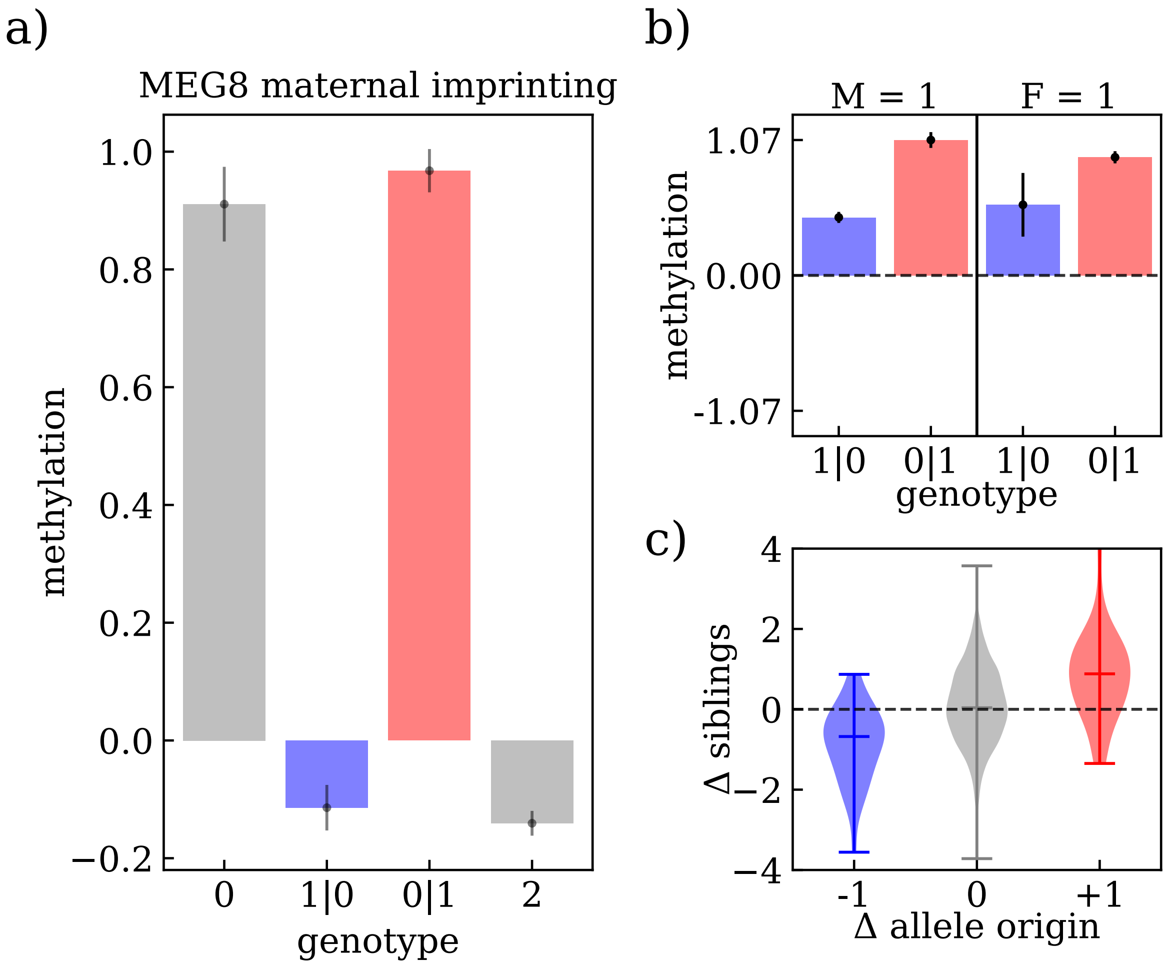


**Figure 4: Methylation patterns for rs10782494 fine–mapped from rs2144819 acting on probe cg06150437, associated with *MEG8*.** (a) Average methylation of all children adjusted for the selected SNP and stratified by genotype, where differences between reciprocal heterozygotes (shown in blue and red) are typically interpreted as evidence of imprinting. (b) Average methylation of reciprocal heterozygotes from heterozygous parents adjusted for the indirect maternal $M$ or paternal $F$ component of the SNP, where similar measurements for both reciprocal heterozygotes from either homozygous mothers or fathers point towards indirect parental effects rather than imprinting. (c) Violin plots of differences in methylation of sibling pairs adjusted for imprinting effect on the SNP, stratified according to parental origin of the alternative allele, where a slope between different sibling pairs indicates an imprinting effect. Error bars in (a) and (b) correspond to the standard deviation.

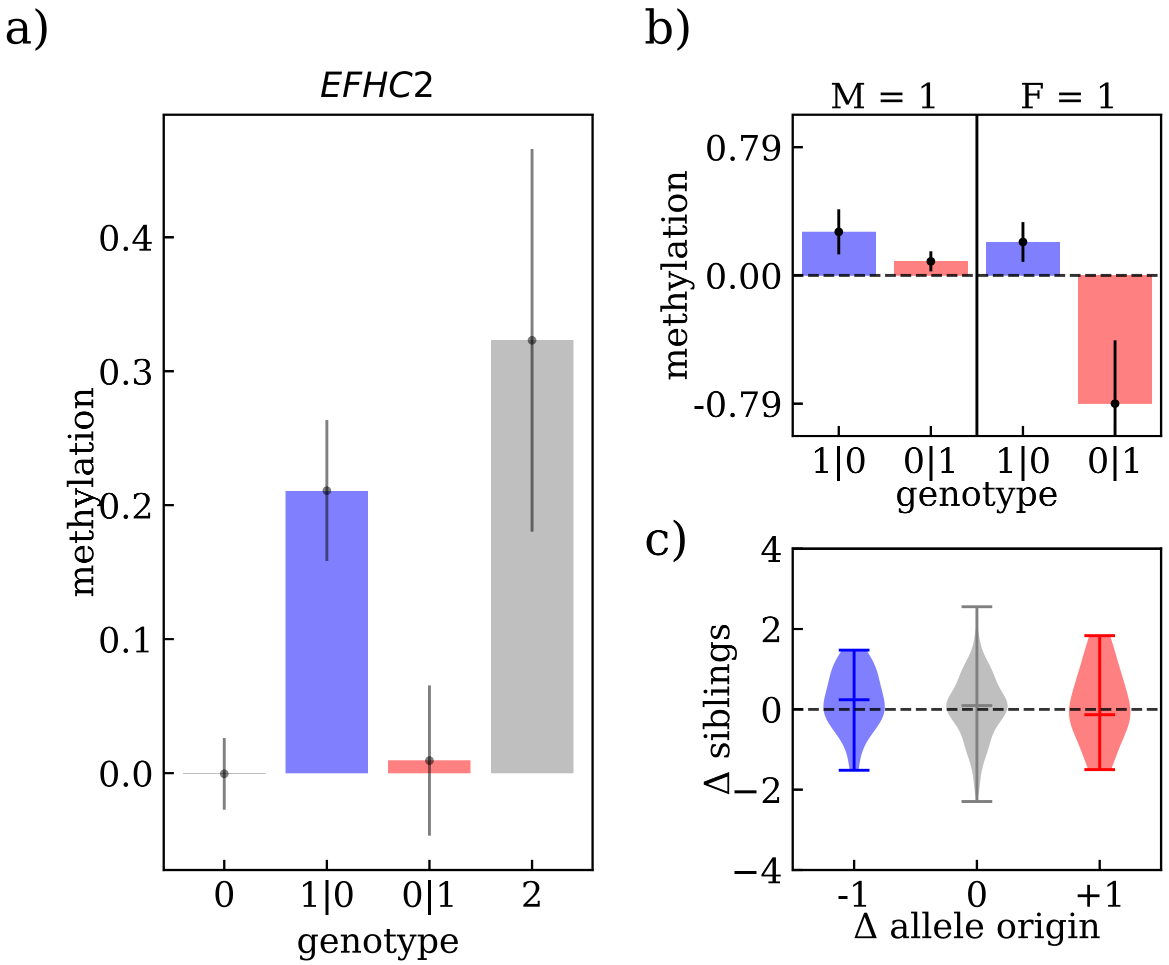


**Figure 5: Methylation patterns for rs7058398 acting on probe cg16088894, associated with *EFHC2*.** (a) Average methylation of all children adjusted for the selected SNP and stratified by genotype, where differences between reciprocal heterozygotes (shown in blue and red) are typically interpreted as evidence of imprinting. (b) Average methylation of reciprocal heterozygotes from heterozygous parents adjusted for the indirect maternal $M$ or paternal $F$ component of the SNP, where similar measurements for both reciprocal heterozygotes from either homozygous mothers or fathers point towards indirect parental effects rather than imprinting. (c) Violin plots of differences in methylation of sibling pairs adjusted for imprinting effect on the SNP, stratified according to parental origin of the alternative allele, where a slope between different sibling pairs indicates an imprinting effect. Error bars in (a) and (b) correspond to the standard deviation.

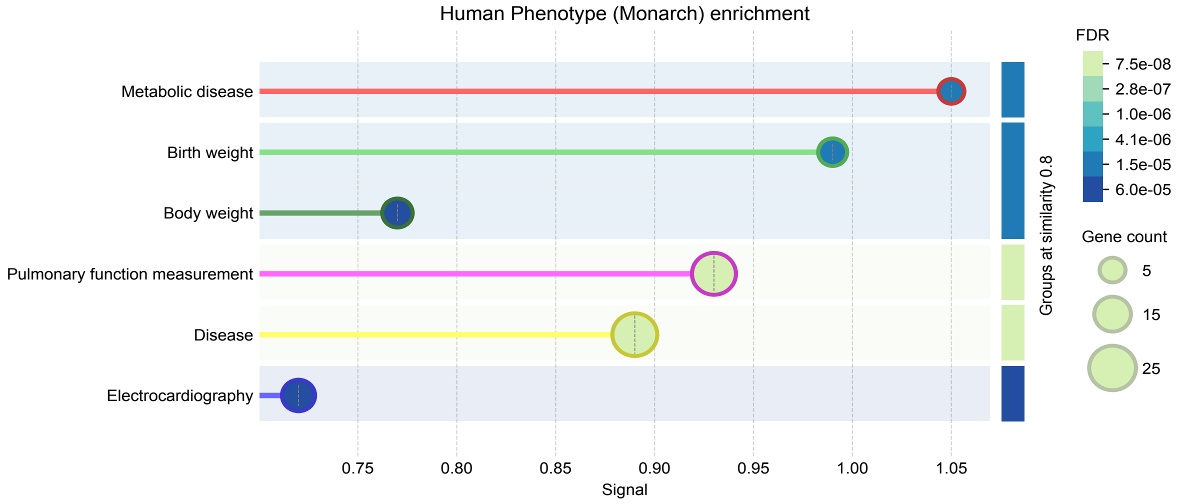


**Figure 6: Phenotype enrichment analysis.** Human phenotype enrichment given by STRING for top 6 traits applied to 68 selected gene identifiers. The phenotypes present are consistent with known functions and associations of imprinted genes.

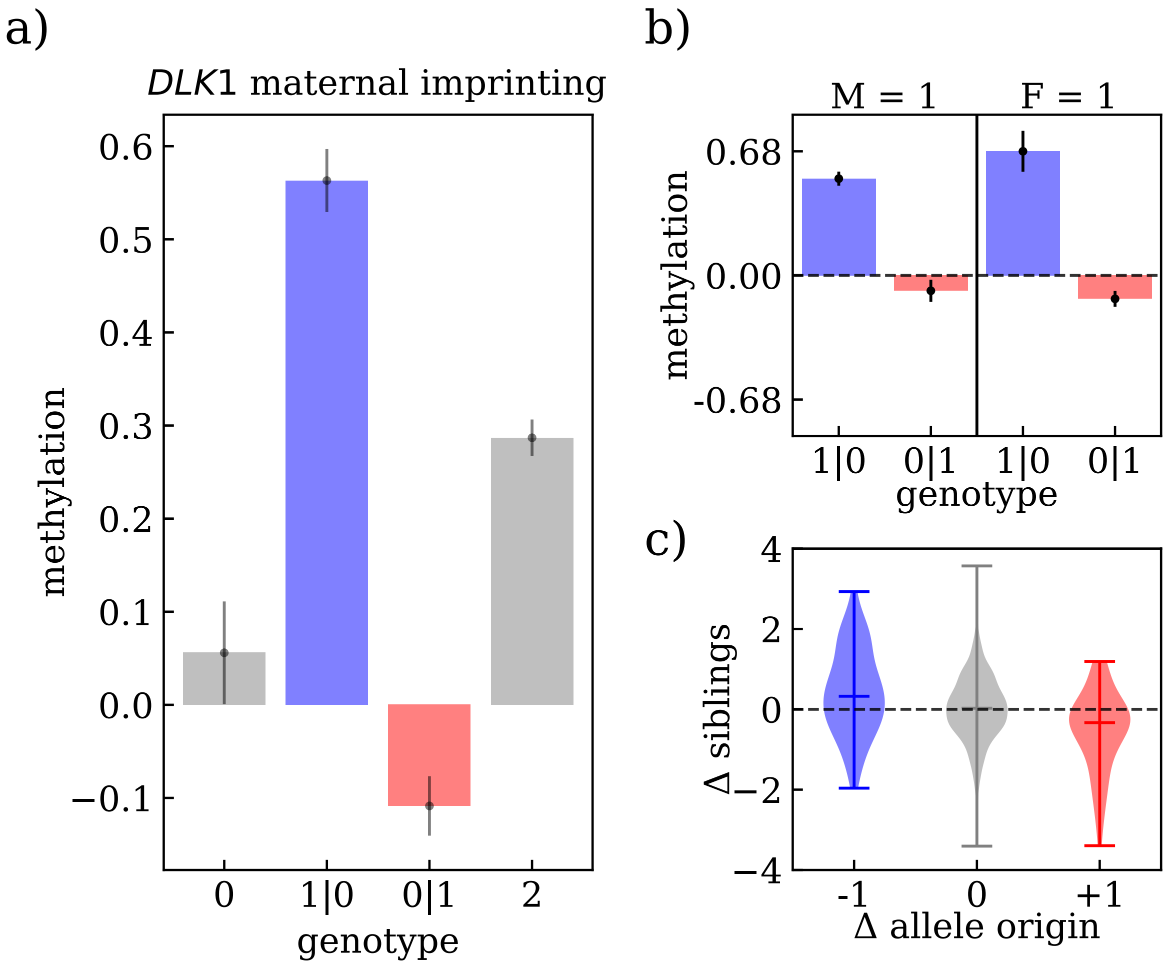


**Figure 7: Methylation patterns for rs12881545 fine–mapped from rs80041283 acting on probe cg18279536, associated with *DLK1*, showing maternal imprinting.** (a) Average methylation of all children adjusted for the selected SNP and stratified by genotype, where differences between reciprocal heterozygotes (shown in blue and red) are typically interpreted as evidence of imprinting. (b) Average methylation of reciprocal heterozygotes from heterozygous parents adjusted for the indirect maternal $M$ or paternal $F$ component of the SNP, where similar measurements for both reciprocal heterozygotes from either homozygous mothers or fathers point towards indirect parental effects rather than imprinting. (c) Violin plots of differences in methylation of sibling pairs adjusted for imprinting effect on the SNP, stratified according to parental origin of the alternative allele, where a slope between different sibling pairs indicates an imprinting effect. Error bars in (a) and (b) correspond to the standard deviation.

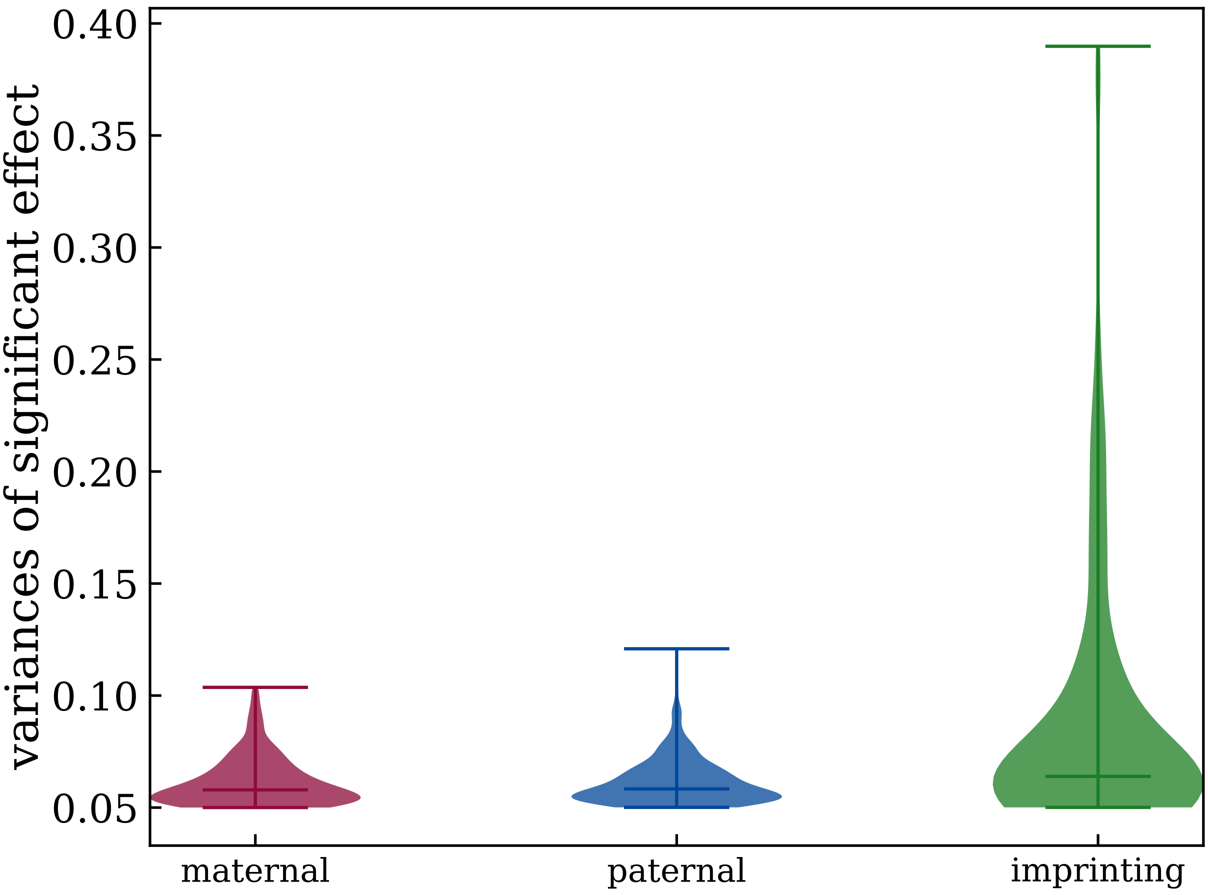


**Figure 8: Violin plots for variances due to indirect maternal (M), indirect paternal (F) and imprinting effects (I) for maternal, paternal and POE–dependent probes.**

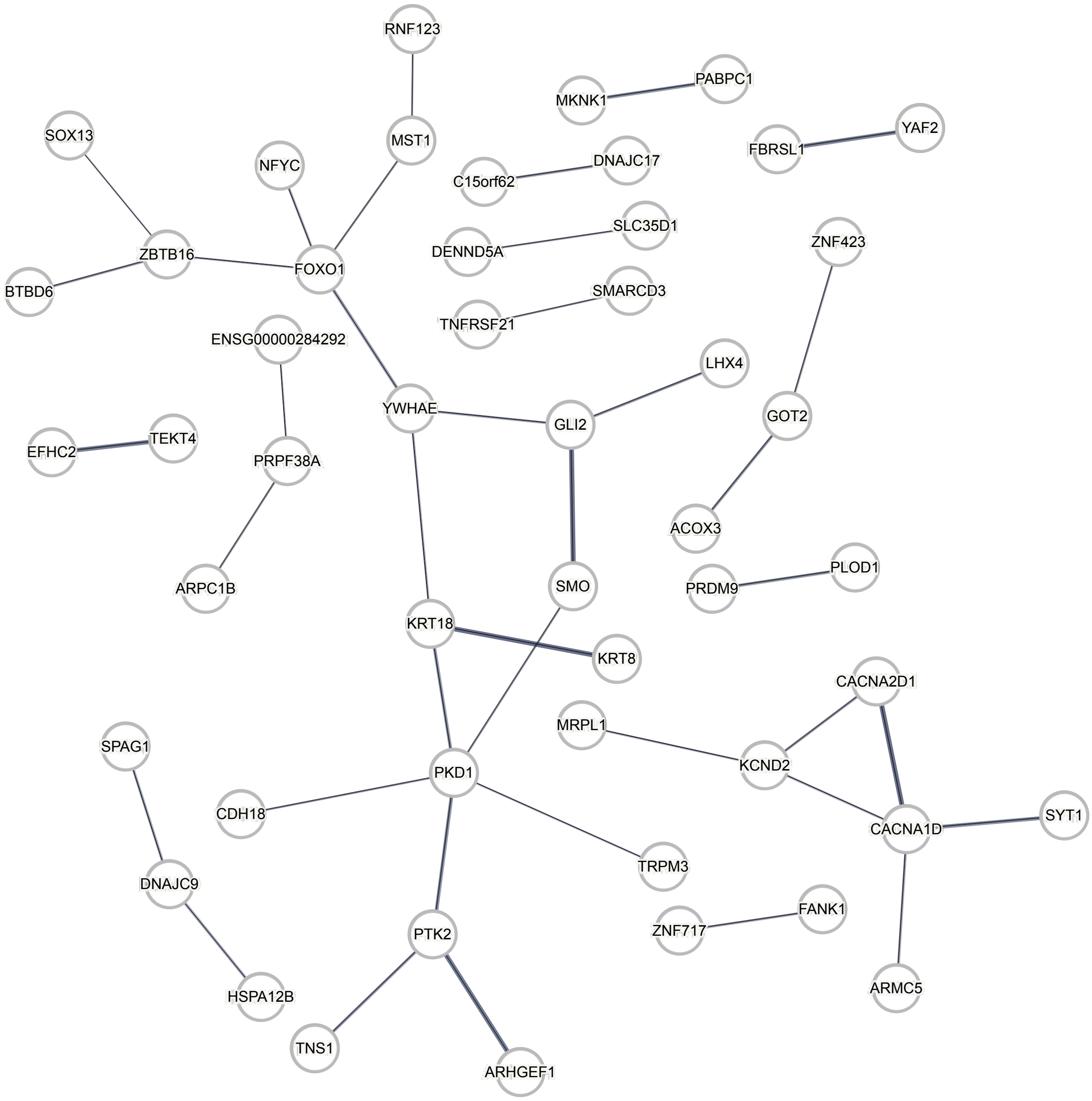


**Figure 9: Protein–protein interaction network for genes associated with indirect maternal dependent probes.** The network is generated using STRING [13] and restricted to 50 genes with interactions, linked to indirect maternal dependent probes.

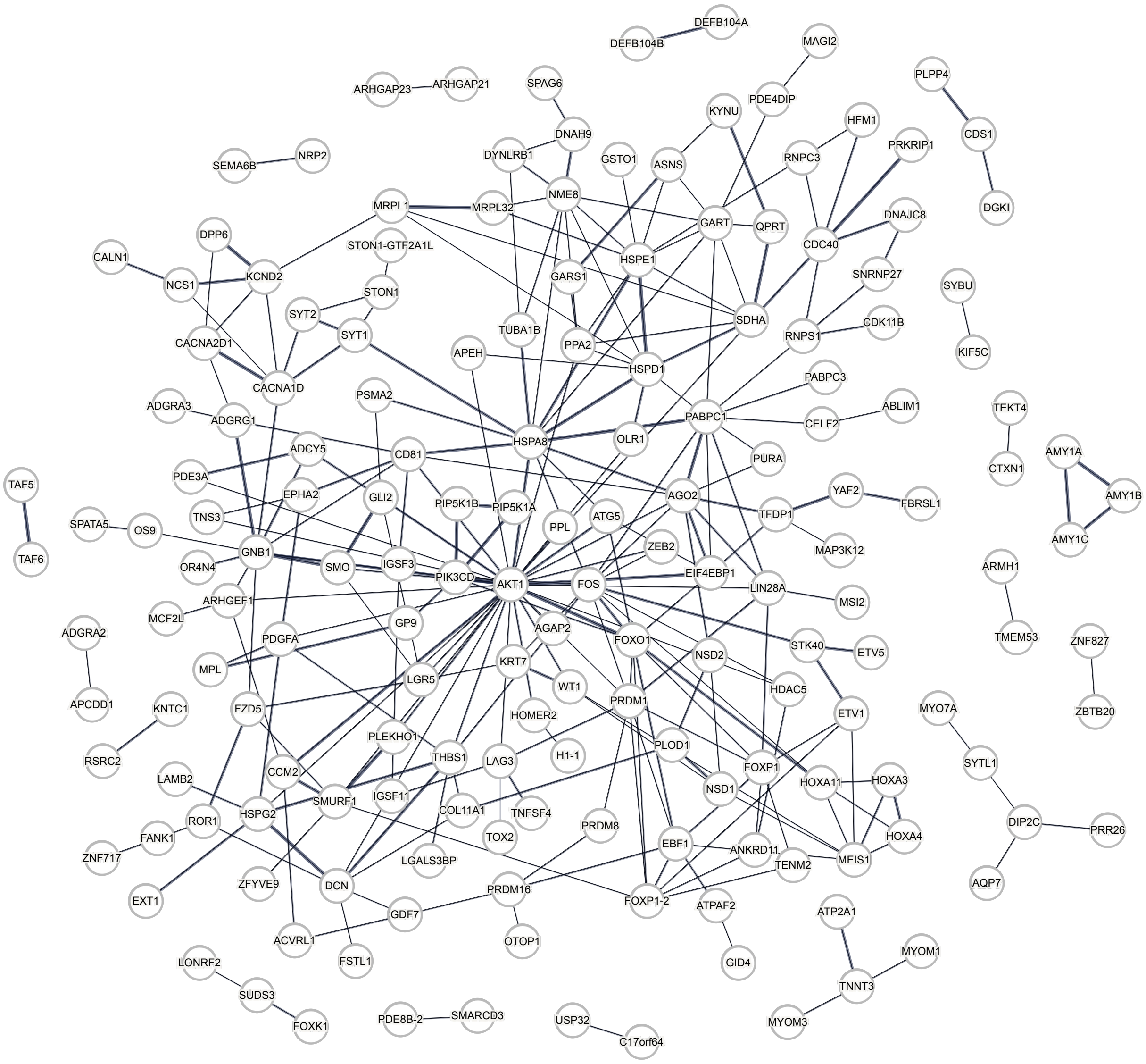


**Figure 10: Protein–protein interaction network for genes associated with indirect paternal dependent probes.** The network is generated using STRING [13] and restricted to 174 genes with interactions, linked to indirect paternal dependent probes.

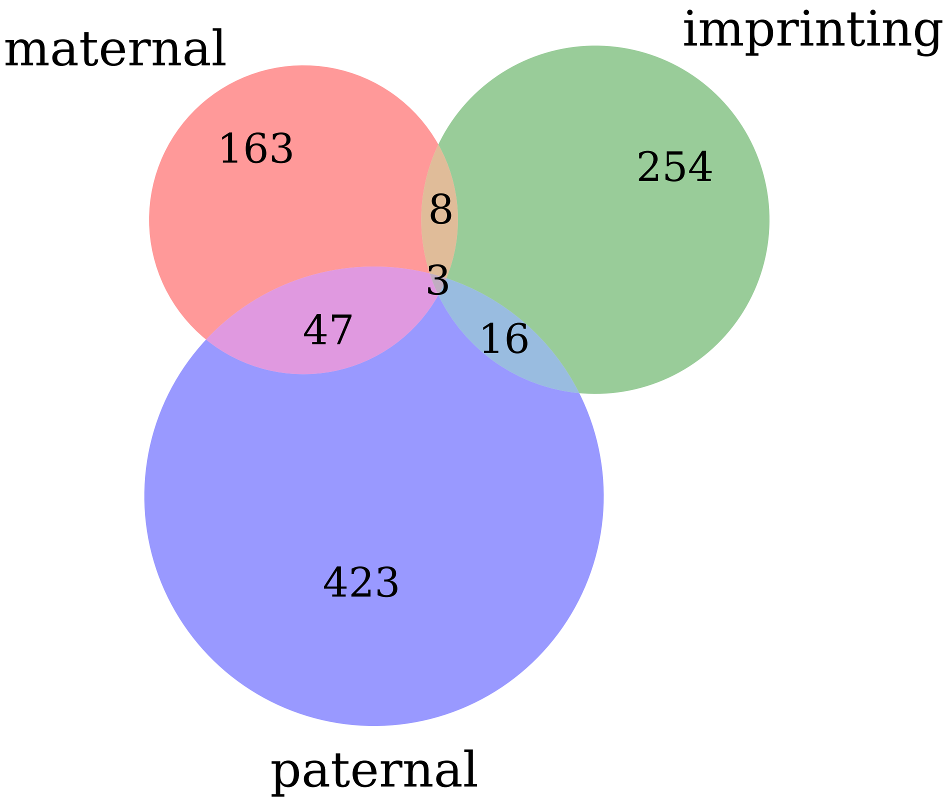


**Figure 11: Venn diagram of identified probes.** Showing the overlap between maternal dependent (red), paternal dependent (blue) and POE–dependent probes (labeled "imprinting", green)

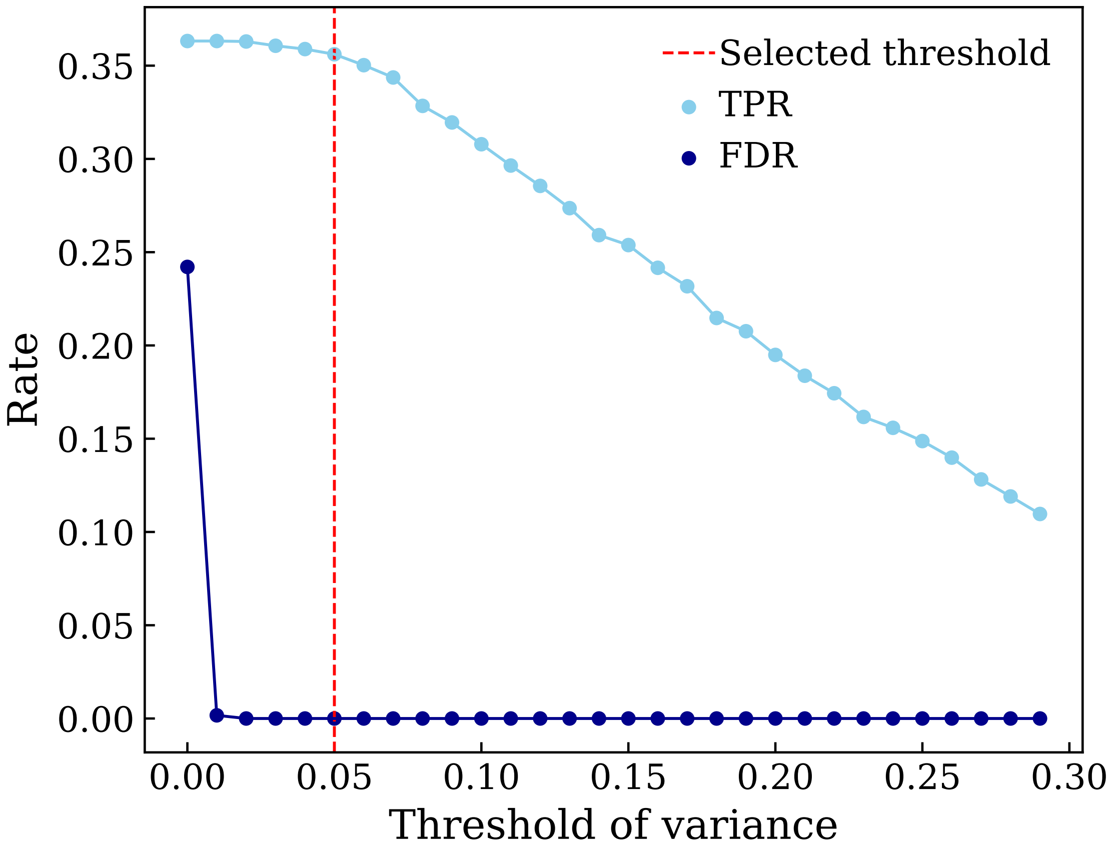


**Figure 12: True positive and false discovery rates as function of the threshold set on the estimated variance using simulated data and the *CMFI* model.** 400 replicates simulating different scenarios of imprinting as well as direct and indirect parental effects are used. The final selected threshold of 0.05 is marked with a dashed red vertical line.

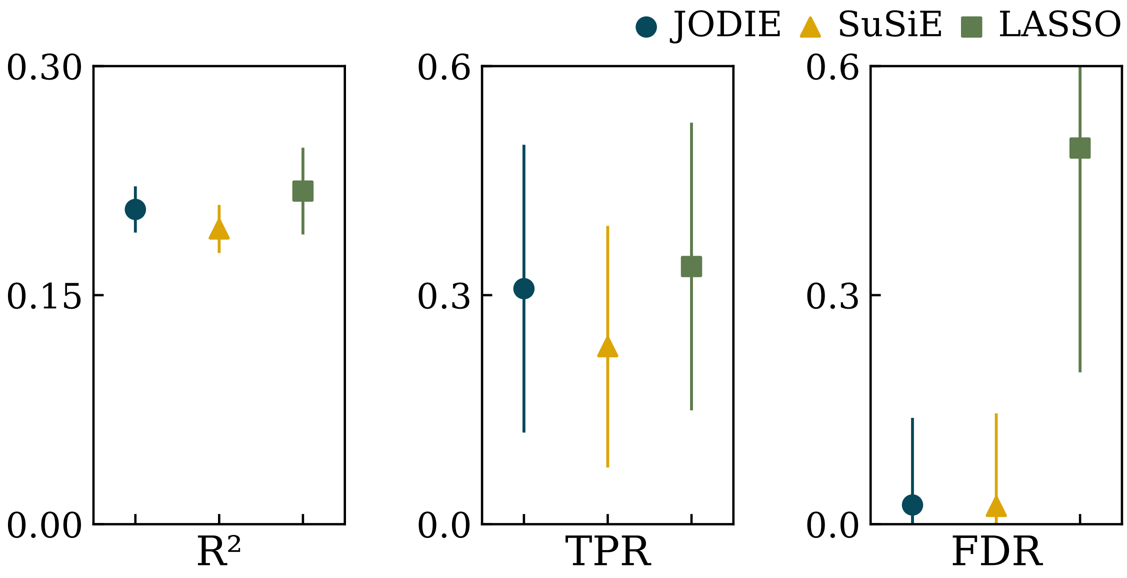


**Figure 13: Performance of the three different regression methods using simulated data.** Squared Pearson correlation coefficient $R^2$ (left), true positive rate TPR (middle), and false discovery rate (FDR) (right) are shown, averaged across 400 replicates simulating different scenarios of imprinting as well as direct and indirect parental effects for regression methods, JODIE (blue), SuSiE (yellow), and LASSO (green). Metrics are calculated based on the correct identification of both the causal SNP and the effect component. The error bars represent the standard deviation.

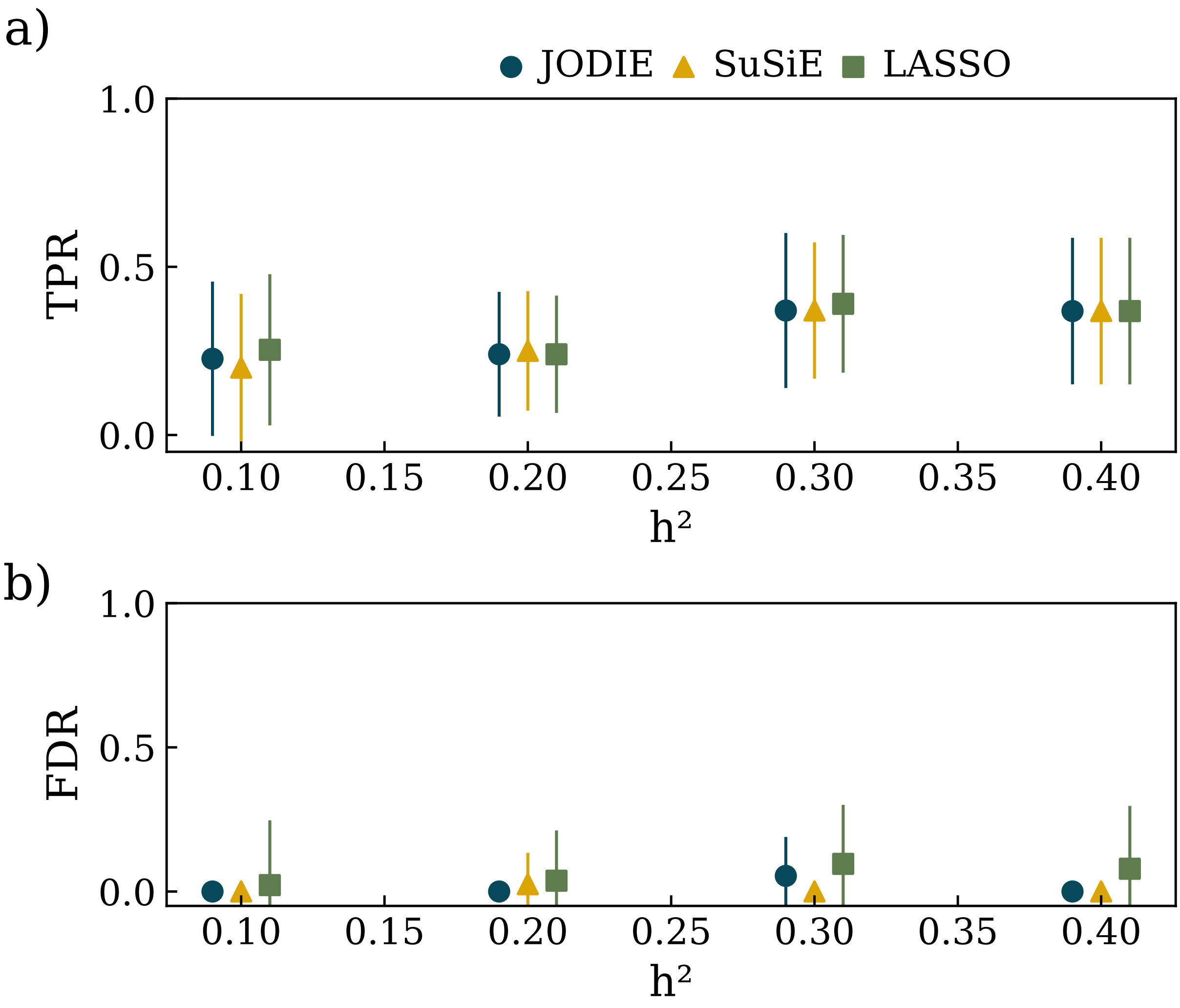


**Figure 14: Performance of the three different regression methods using simulated data with different heritability values.** True positive rate TPR (a) and false discovery rate FDR (b) are shown as function of $V = h^2$, averaged across 20 replicates, simulating only direct effects with different $V = h^2$ values for the three regression methods JODIE (blue), SuSiE (yellow), and LASSO (green). The error bars represent the standard deviation.

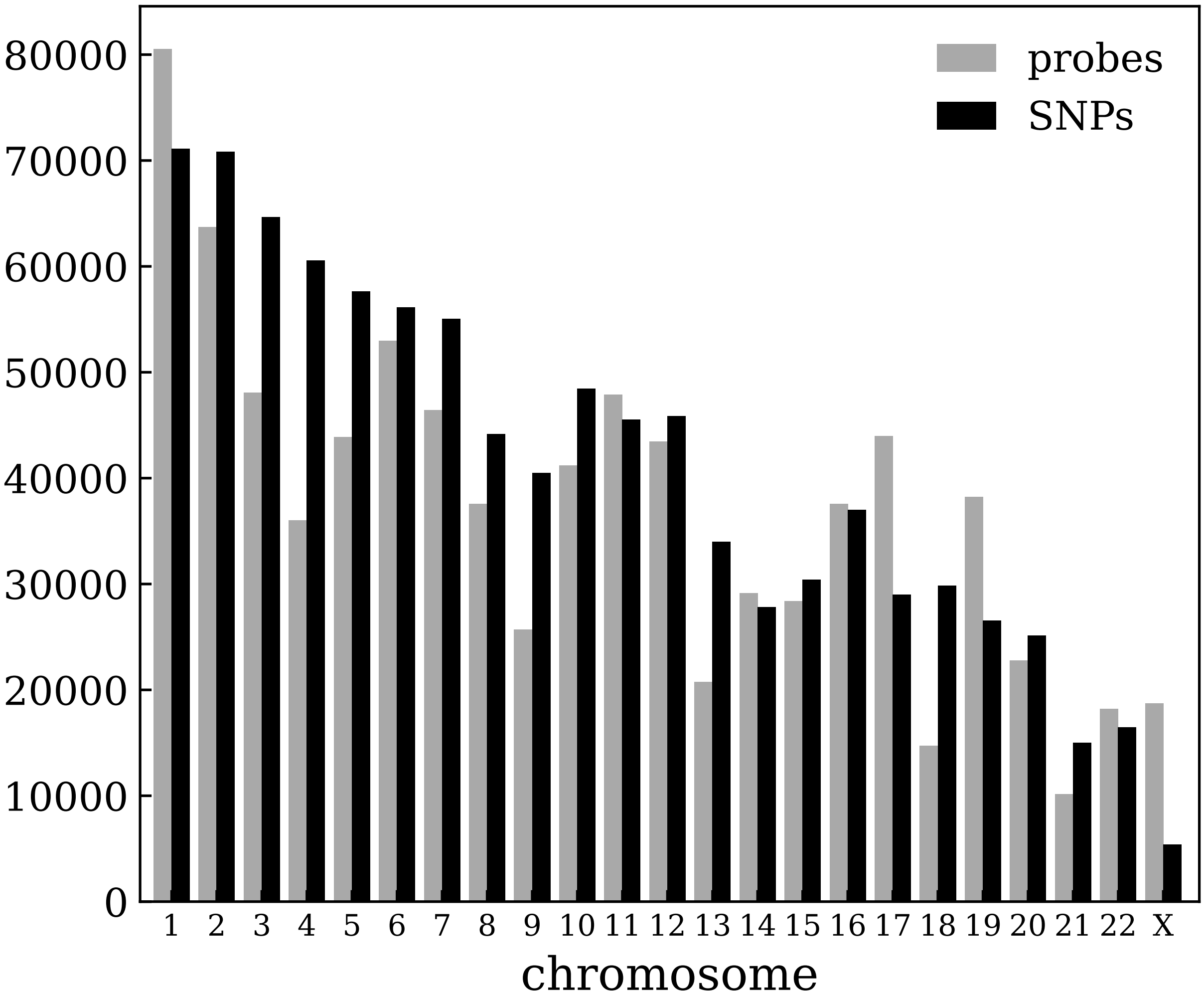


**Figure 15: Number of methylation probes and SNPs per chromosome.** Number of total methylation probes (grey) per chromosome in the Generation Scotland data analyzed in this study and SNPs (black) included in the regression model for the chromosome the probe is located on.

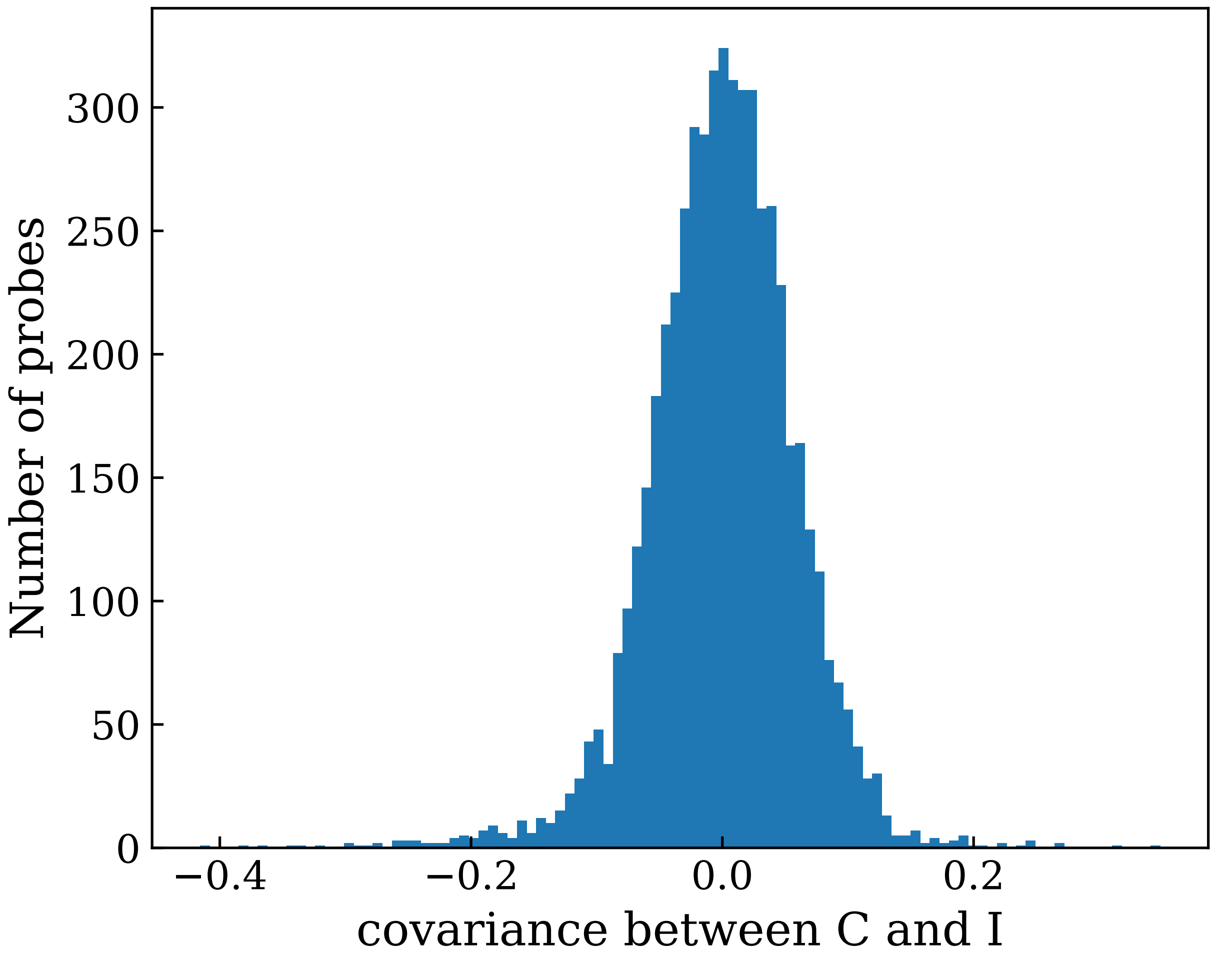


**Figure 16: Histogram of the estimated covariance between direct effects and POEs for the 5,435 probes initially selected with the *CI* model.** The covariance between direct effects ($C$) and POEs ($I$) allows us to determine the type of imprinting. A negative value corresponds to maternal imprinting/paternal expression while a positive one indicates paternal imprinting/maternal expression. The negative side of the distribution is more voluminous, corresponding to more probes showing apparent paternal expression.

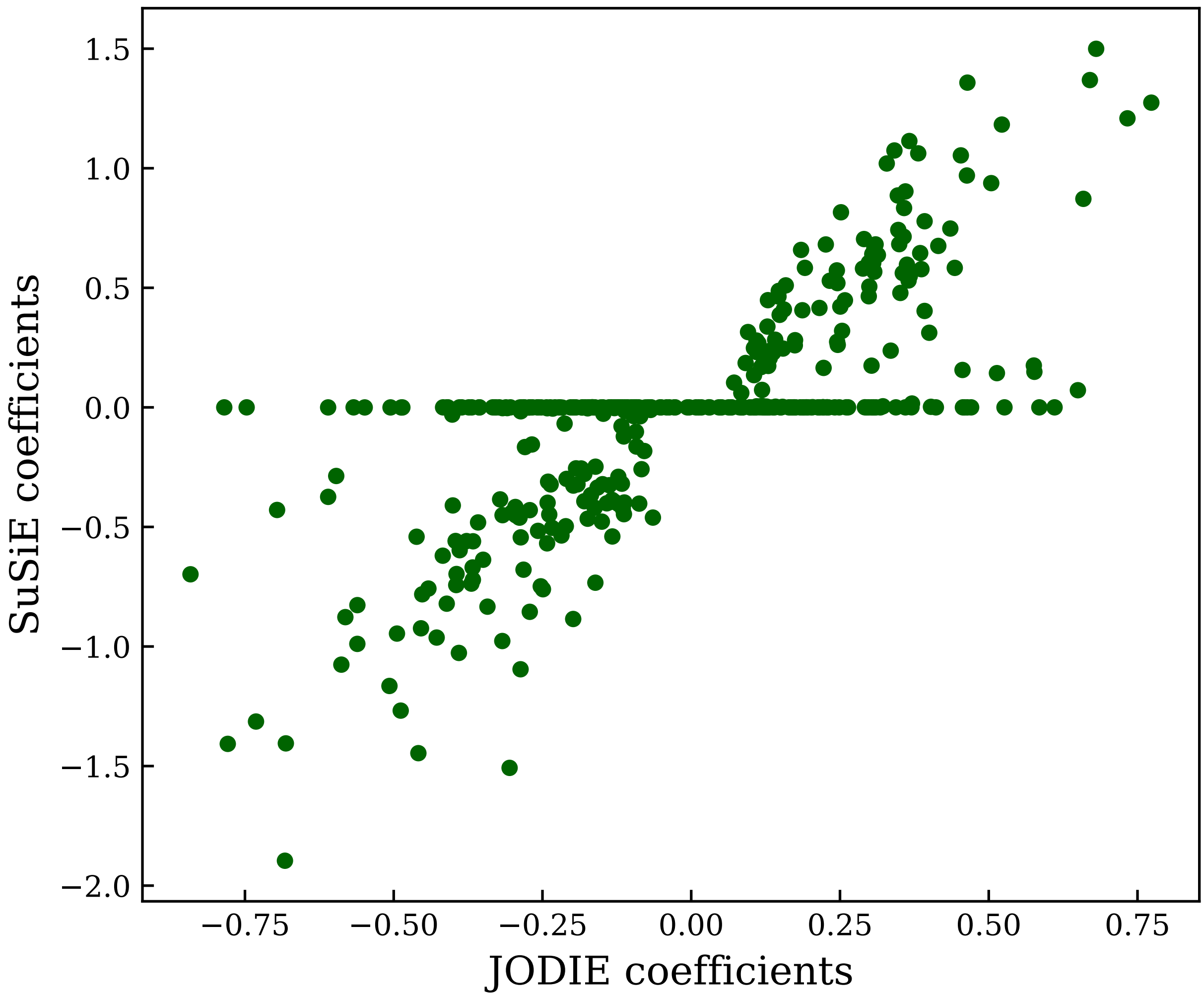


**Figure 17: Estimated POEs using the *CMFI* model and SuSiE for 412 identified SNPs associated with POE–dependent probes.** SuSiE is applied to a reduced set of cis–SNPs ($< 5$Mb) which are shown here.

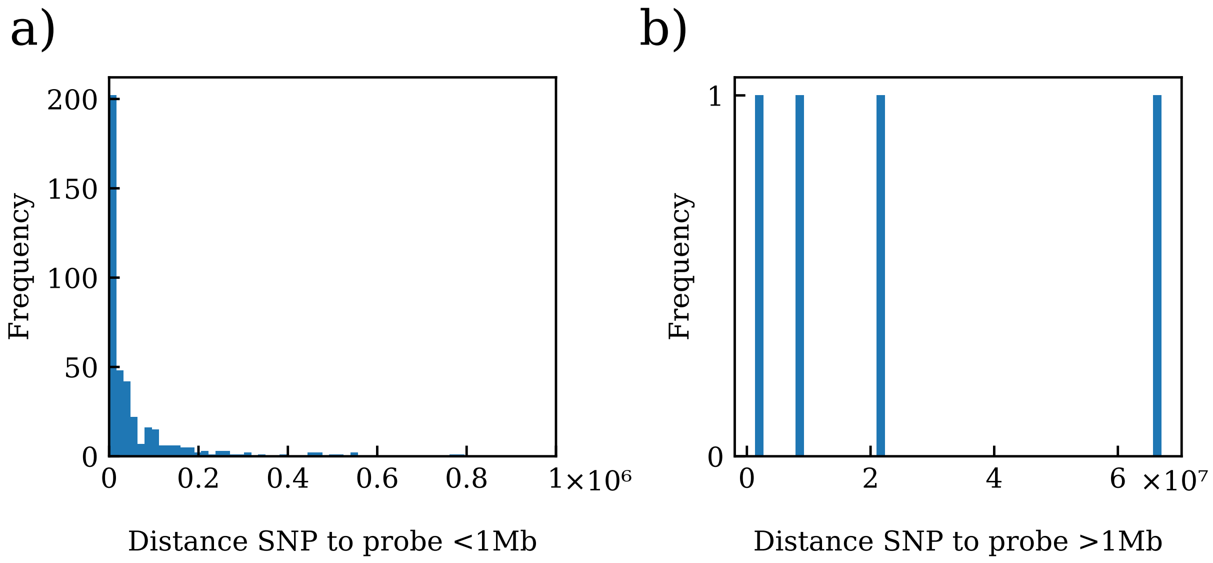


**Figure 18: Distances between POE–dependent probes and associated SNPs.** (a) Histogram of distances of probes and associated cis ($< 1$Mb). (b) Histogram of distances of probes and associated trans ($> 1$Mb).

#### 2 Supplementary Datasets

1. **SD_POEprobes.xlsx**Identified POE–dependent probes. Chromosome, probe ID, position, calculated variances and covariances using JODIE and $CMFI$ model, gene associations given by Generation Scotland annotation file.

2. **SD_fineMapped.xlsx** Fine–mapped SNPs associated with POE–dependent probes. In the case of the same fine–mapped SNP being associated with the same probe through different windows (original SNPs) the number of iterations were summed, variances and coefficients are calculated as an average and values referring to multiple windows are separated by ";". Columns paternal, maternal, complex, direct, indirect maternal and indirect paternal refer to the number of iterations where the SNP presents each imprinting pattern and effect respectively. Iterations column shows the number of total times a SNP is selected for the regression for that probe. Credible set is an identifier for the number of the SNP group to which the fine–mapped SNP is part of, separated by ";" in the case of multiple windows. Gene probe given by Generation Scotland annotation file.

3. **SD_genes.xlsx** Gene annotation for probes and associated SNPs. Sheet 1 SD_POEgenes: Gene annotation of POE–dependent probes. Sheet 2 SD_SNPgenes: Gene annotation of POE–dependent associated SNPs. Sheet 3 SD_maternalGenes: Gene annotation of indirect maternal dependent probes. Sheet 4 SD_paternalGenes: Gene annotation of indirect paternal dependent probes

4. **SD_indirectProbes.xlsx** Sheet 1 SD_maternalProbes: Identified maternal dependent probes, regression results using JODIE and $CMFI$ model. Sheet 2 SD_paternalProbes: Identified paternal dependent probes, regression results using JODIE and $CMFI$ model.

#### References


[1] Fei Yang, Zhuang Xiong, Wenting Zong, Demian Kong, Bixia Tang, Xupeng Chen, Yaoke Wei, Xiangyu Yu, Yiran Zhang, Dong Zou, Zhang Zhang, Yiming Bao, and Rujiao Li. EWAS open platform 2026: a deeply integrated resource for epigenome-wide association studies. *Nucleic Acids Res.*, 54(D1):D1061–D1068, January 2026.

[2] Zhuang Xiong, Fei Yang, Mengwei Li, Yingke Ma, Wei Zhao, Guoliang Wang, Zhaohua Li, Xinchang Zheng, Dong Zou, Wenting Zong, Hongen Kang, Yaokai Jia, Rujiao Li, Zhang Zhang, and Yiming Bao. EWAS open platform: integrated data, knowledge and toolkit for epigenome-wide association study. *Nucleic Acids Res.*, 50(D1):D1004–D1009, January 2022.

[11] Morgan E Levine, Ake T Lu, Austin Quach, Brian H Chen, Themistocles L Assimes, Stefania Bandinelli, Lifang Hou, Andrea A Baccarelli, James D Stewart, Yun Li, Eric A Whitsel, James G Wilson, Alex P Reiner, Abraham Aviv, Kurt Lohman, Yongmei Liu, Luigi Ferrucci,

and Steve Horvath. An epigenetic biomarker of aging for lifespan and healthspan. *Aging (Albany NY)*, 10(4):573–591, April 2018.

[12] Flavio Santilli and Ana Boskovic. Mechanisms of transgenerational epigenetic inheritance: lessons from animal model organisms. *Curr. Opin. Genet. Dev.*, 79(102024):102024, April 2023.

[13] Damian Szklarczyk, Rebecca Kirsch, Mikaela Koutrouli, Katerina Nastou, Farrokh Mehryary, Radja Hachilif, Annika L Gable, Tao Fang, Nadezhda T Doncheva, Sampo Pyysalo, Peer Bork, Lars J Jensen, and Christian von Mering. The string database in 2023: protein–protein association networks and functional enrichment analyses for any sequenced genome of interest. *Nucleic Acids Research*, 51(D1):D638–D646, November 2022.

[23] Ilse Krätschmer, Laura Hegemann, Gunnar Pálsson, Robin Hofmeister, Anne Richmond, Caroline Hayward, Ole A. Andreassen, Astanand Jugessur, Ida Elken Sønderby, Adrian Dahl Askelund, Estonian Biobank Research Team, Zoltan Kutalik, Riccardo E. Marioni, Marteinn Thor Hardarson, Ólafur A. Stefansson, Hákon Jónsson, Peter Kharchenko, Kári Stefánsson, Bjarni V. Halldórsson, Alexandra Havdahl, and Matthew R. Robinson. Sex differences in the genetic basis of human recombination within 190,000 parent-child pairs. *bioRxiv*, 2026. Preprint posted May 2, 2026.
